# Using Parsimony-Guided Tree Proposals to Accelerate Convergence in Bayesian Phylogenetic Inference

**DOI:** 10.1101/778571

**Authors:** Chi Zhang, John P. Huelsenbeck, Fredrik Ronquist

## Abstract

Sampling across tree space is one of the major challenges in Bayesian phylogenetic inference using Markov chain Monte Carlo (MCMC) algorithms. Standard MCMC tree moves consider small random perturbations of the topology, and select from candidate trees at random or based on the distance between the old and new topologies. MCMC algorithms using such moves tend to get trapped in tree space, making them slow in finding the globally most probable trees (known as ‘convergence’) and in estimating the correct proportions of the different types of them (known as ‘mixing’). Here, we introduce a new class of moves, which propose trees based on their parsimony scores. The proposal distribution derived from the parsimony scores is a quickly computable albeit rough approximation of the conditional posterior distribution over candidate trees. We demonstrate with simulations that parsimony-guided moves correctly sample the uniform distribution of topologies from the prior. We then evaluate their performance against standard moves using six challenging empirical datasets, for which we were able to obtain accurate reference estimates of the posterior using long MCMC runs, a mix of topology proposals, and Metropolis coupling. On these datasets, ranging in size from 357 to 934 taxa and from 1,740 to 5,681 sites, we find that single chains using parsimony-guided moves usually converge an order of magnitude faster than chains using standard moves. They also exhibit better mixing, that is, they cover the most probable trees more quickly. Our results show that tree moves based on quick and dirty estimates of the posterior probability can significantly outperform standard moves. Future research will have to show to what extent the performance of such moves can be improved further by finding better ways of approximating the posterior probability, taking the trade-off between accuracy and speed into account.

## Introduction

Introduced to the field two decades ago (Rannala and Yang, 1996; Yang and Rannala, 1997; Mau and Newton, 1997; Li et al., 2000), Bayesian estimation of phylogeny has become widely used by evolutionary biologists (see Huelsenbeck et al., 2001; Holder and Lewis, 2003; Yang and Rannala, 2012; Yang, 2014; Nascimento et al., 2017, for reviews). The Bayesian framework is attractive for many reasons, including the simple interpretation of results, the ability to address interesting evolutionary questions, and the availability of efficient and easy-to-use computer programs that implement it (Ronquist and Huelsenbeck, 2003; Ronquist et al., 2012b; Drummond and Rambaut, 2007; Bouckaert et al., 2014; Höhna et al., 2016).

Bayesian estimation of phylogeny almost always relies on Markov chain Monte Carlo (MCMC: Metropolis et al., 1953; Hastings, 1970) to sample trees in proportion to their posterior probabilities (for exceptions, see Bouchard-Côté et al., 2012; Wang et al., 2016). The MCMC procedure typically uses a mixture of different proposal mechanisms (also called moves or operators) that change one or a few of the parameters in the model. Unfortunately, devising MCMC proposals that sample well across the space of evolutionary trees is quite challenging. For any reasonable number of tips, a MCMC proposal has a huge space of tree topologies to explore. It also needs to tackle the complex dependencies between topology and branch lengths. Most MCMC tree proposals studied and used in current Bayesian MCMC phylogenetic software are stochastic versions of tree-perturbation methods that were originally used to find optimal parsimony or maximum likelihood trees in hill-climbing algorithms. The basic perturbation methods include nearest neighbor interchange (NNI), subtree pruning and regrafting (SPR), and (for non-clock trees) tree bisection and reconnection (TBR) (Larget and Simon, 1999; Lakner et al., 2008). Each of these mechanisms can generate a set of candidate trees from the current one by applying a particular type of tree modification.

In a hill-climbing algorithm, one would typically score all candidate trees generated by one of these mechanisms, and then choose the best one. In the MCMC context, a new tree is instead drawn from a suitable probability distribution over the candidate trees. The simplest choice is a uniform distribution, and this is still a common choice for stochastic NNI (sNNI) moves. Uniformly random SPR and TBR proposals, however, are inefficient because they tend to make such drastic changes that the new tree has negligible posterior probability and will be rejected. A better option is to bias SPR and TBR moves towards more modest changes as in the “extending” SPR (eSPR) and “extending” TBR (eTBR) moves introduced in MrBayes (Huelsenbeck and Ronquist, 2001; Ronquist and Huelsenbeck, 2003). These proposals move the regrafting point away from the pruning point in a step-wise fashion, applying a constant probability in each step to decide whether the distance should be further extended. sNNI, eSPR and eTBR are among the most efficient MCMC tree moves known for non-clock trees (Lakner et al., 2008), but given the frequent observation of empirical tree spaces that fail to yield to these proposals, even when Metropolis-coupling (Geyer, 1991) is used, there is clearly room for improvement.

One possibility would be to choose among candidate trees according to their posterior probability. This is known as a ‘Gibbs’ proposal (Liu, 2004), and it will always be accepted. The mixing rate of a Gibbs proposal can be further improved by ‘Metropolizing’ the move, that is, by removing the probability of proposing the starting state as the new state (Peskun, 1973). Unfortunately, it is computationally costly to obtain the posterior probability of any set of alternative trees that is reasonably large. Therefore, Gibbs or Metropolized Gibbs proposals are quite difficult to implement except in some very unusual circumstances (Huelsenbeck et al., 2008). Gibbs sampling is a special case of the Metropolis-Hastings algorithm. In general, the proposal ratio (also known as the “Hastings ratio”) of the Metropolis-Hastings algorithm is used to adjust the acceptance ratio, correcting for any asymmetry that may arise between forward and backward proposals.

Instead of using posterior probabilities directly, we could guide tree proposals based on an approximation of the posterior distribution over topologies obtained in a previous analysis (Höhna and Drummond, 2012). Such approximations are best computed from the sampled split (taxon bipartition) frequencies, either alone or in combinations of two or three adjacent splits (Ronquist et al., 2004; Höhna and Drummond, 2012; Larget, 2013). Guiding tree proposals based on such approximations can significantly improve their performance (Höhna and Drummond, 2012). A potential concern with this approach is that the efficiency of the final run could be affected by any errors or biases in the approximation of the posterior distribution over topologies obtained in the preliminary run. In other words, this approach pushes the most difficult challenge for tree proposals to the preliminary analysis, when an approximation of tree space is not available.

In this paper, we explore another idea, namely to guide tree proposals based on a quick and dirty approximation of the posterior probability of trees, computed on the fly. Specifically, we base the approximation on the parsimony score of topologies, inspired by the links that do exist between parsimony scores and probability (Huelsenbeck et al., 2008). To our knowledge, such parsimony-guided tree proposals were first introduced in MrBayes 3.2 (Ronquist et al., 2012b), where they were included in the default set of tree moves based on promising preliminary experiments. In the development of ExaBayes (Aberer et al., 2014), parsimony-guided tree proposals modeled after MrBayes were found to be essential for convergence on large datasets (A. Aberer, pers. comm.). However, these proposals have never been described in the literature, nor have their properties been examined in detail.

Here, we describe a couple of variants of parsimony-guided SPR (pSPR) and parsimony-guided TBR (pTBR) proposals, which have been implemented in MrBayes. We use simulations to verify the implementations, and we use analyses of empirical datasets to show that pSPR and pTBR moves significantly outperform eSPR and eTBR both in terms of convergence and mixing. Tree moves have received surprising little attention, given their importance in Bayesian MCMC phylogenetics (for exceptions, see Lakner et al., 2008; Höhna et al., 2008; Höhna and Drummond, 2012; Whidden and Matsen, 2015). We hope that this paper will stimulate further work on guided MCMC proposals and other ideas for improving MCMC exploration of tree space.

## Methods

In this section, we first describe the tree proposals used in the paper. We then show the simulations we used to verify the implementation of the parsimony-guided proposals. Finally, we describe the empirical tests we used to evaluate the performance of the parsimony-guided tree proposals against standard extending tree proposals.

All algorithms were implemented in MrBayes version 3.2.7, available from GitHub (https://github.com/NBISweden/MrBayes). The MrBayes scripts used in the analyses are available as Supplementary Material. We focus the description here on the tree proposals; details of the other proposals and prior settings used in the analyses are available in the MrBayes scripts and in the manual.

### Tree Proposals

We focus entirely on tree proposals for unrooted trees. Specifically, we explore two variants each of pSPR and pTBR, and compare their performance to eSPR and eTBR. In the reference runs, we also use sNNI. For a more complete description of the MCMC context of these proposals, see Appendix; here we focus only on the essential details needed to describe them unambiguously. Note that the eSPR move used here is different from that examined in a previous study (Lakner et al., 2008) in that the initial branch is picked from all branches in the move, rather than only from internal branches.

### Nearest Neighbor Interchange (NNI)

First, an internal branch is picked at random. Label the branches with subtrees that are incident to the chosen branch A, B, C and D, such that the original topology is ((A,B),C,D). Without loss of generality, assume that D stays in its original position. Then the topology is modified by exchanging A and C, or B and C. In the stochastic Metropolized version of the move, denoted sNNI here, the two alternative topologies are chosen with equal probability. The move leaves the topology with the same taxon bipartitions except for the bipartition associated with the initially chosen internal branch. This gives a natural mapping of branches between the trees. The branch lengths of the old tree are applied unmodified to the new tree using this mapping. The proposal ratio is 1.0.

### Subtree Pruning and Regrafting (SPR)

The SPR move first picks a branch *a* (internal or external, with length *υ_a_*) at random. It then randomly selects one of the two subtrees incident to the branch; label this subtree A (Fig. 1a). If *a* is a terminal branch, A will be the tip node. A branch *r* in the remaining subtree, with length *υ_r_*, is chosen for regrafting of A (Fig. 1a). The move is Metropolized, such that a topology change is guaranteed. This means that branch *b* or *q* cannot be selected as *r*. The exact mechanism for choosing *r* varies depending on the type of SPR move (see below). The branch connecting *a* in the moving direction (that is, the branch closer to *r*, *q* in this case, with length *υ_q_*) is also pruned away and moved together with A, leaving the remaining subtree BCDE (Fig. 1b). Subtree A with branches *a* and *q* is inserted before branch *r* connecting subtree E, resulting in *r* connecting *a* and *q* (Fig. 1c). Note that NNI is a special case of SPR: if A is moved one node away to C (*c* is chosen for regrafting) or DE (*p* is chosen for regrafting), this is equivalent to a NNI move around branch *q*.

**Figure 1:**
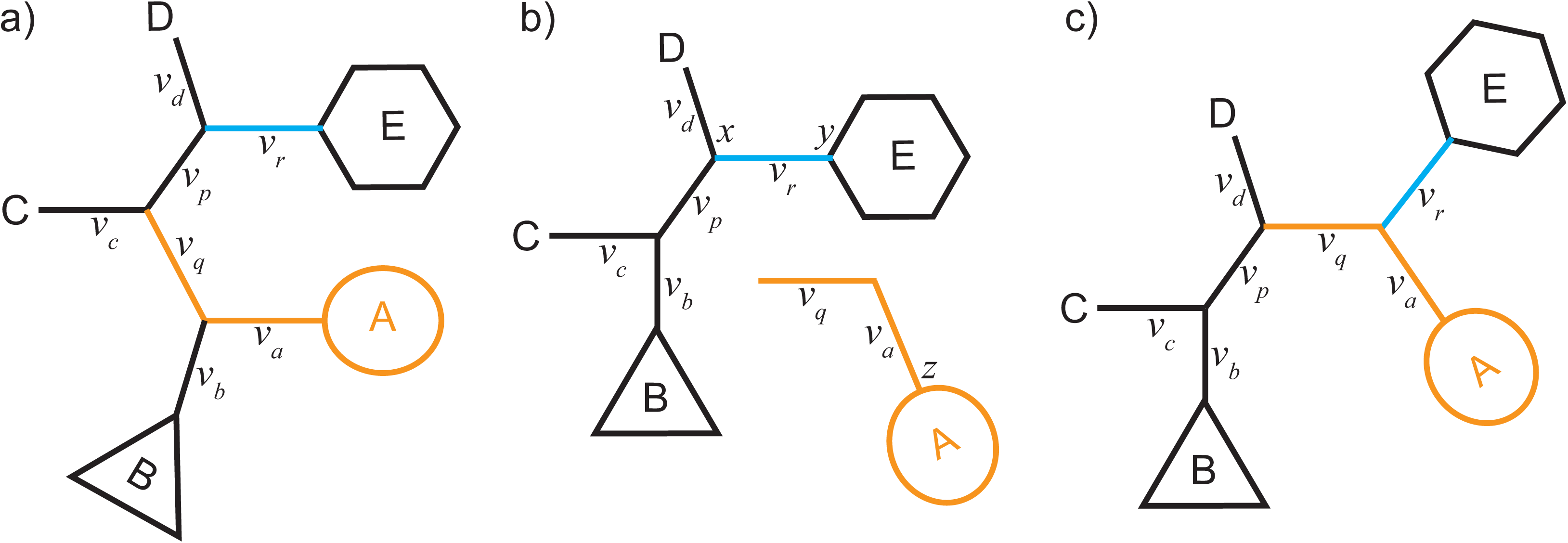
The basic logic of the SPR proposal mechanisms. First, a branch *a* (with length *υ_a_*) is picked at random from all branches in the tree. One of the subtrees incident to *a* is then picked at random and pruned away; this subtree is labeled A here. For regrafting, a branch *r* is picked in the other subtree (labeled BCDE). The branch *q* in the moving direction (the pendant branch) is moved together with *A*, and inserted adjacent to *r*, resulting in *r* connecting *a* and *q*. The specific SPR variants differ in how they pick *r*.

The eSPR move applies an extension mechanism to choose *r*: with probability 1/2, it moves the regrafting point one branch away in either direction (that is, into the B or the CDE subtree). Then with probability *p_e_* (the extension probability), the regrafting point is moved one branch further, and with probability (1 − *p_e_*) it stays at the current location (Fig. 6 in Lakner et al., 2008). If the branch is moved one branch further away, one of the two possible directions is chosen with probability 1/2, and the cycle is repeated. If the extension mechanism encounters a tip, it stops. We refer to a proposal that encounters a tip as a “constrained” proposal. The proposal ratio is 1.0 if the extension mechanism is unconstrained or constrained in both directions of the move. If the move is constrained in just one direction, the proposal ratio becomes 1/(2(1 − *p_e_*)) (backward move constrained) or 2(1 − *p_e_*) (forward move constrained) (Lakner et al., 2008).

The pSPR moves use a mechanism based on parsimony scores to choose *r*. Let *B* be the set of all branches in subtree BCDE, that is, the tree remaining after subtree A is pruned away (Fig. 1b). The weight for proposing branch *i* for regrafting of A, *ω_i_*, is given by:

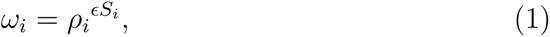

where *ϵ* is the warp factor, *ρ_i_* is the base factor, and *S_i_* is the parsimony score at branch *i*. For computational convenience (see Appendix), *S_i_* at branch *i* is the parsimony score of the tree after regrafting A at *i*, minus the sum of the scores of the two subtrees A and BCDE. We call the tuning parameter *ϵ* the warp factor because it determines how much the parsimony score will influence the weight, that is, the extent to which a uniform probability distribution over candidate trees is modified by the parsimony score. The larger the warp factor, the more heavily the parsimony score will influence the probability of candidate trees being proposed.

Assume that in the set *B*, the branch at the pruning point is *b* and the branch at the regrafting point is *r*. Then the proposal ratio of a pSPR move is

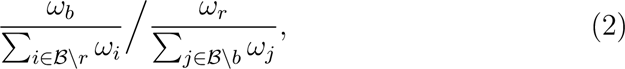

where *B*\*i* is the set of all branches in *B* except *i*. The weights are constructed such that candidate trees with lower (better) parsimony scores receive higher weights and are proposed more often. The proposal ratio corrects for this preference: a proposed move to a more parsimonious tree is likely to be accepted only if the proposal bias is matched by a similar or larger increase in likelihood. Thus, a correctly implemented parsimony-guided proposal will not be affected by parsimony artefacts, such as long-branch attraction.

Here, we employ two schemes for converting parsimony scores to proposal weights. In the first scheme, pSPR1, we simply set all *ρ_i_* to *e*^−1^, such that the weight *ω_i_* = *e*^*−ϵ*_1_*S_i_*^. In the second scheme, pSPR2, we try to accommodate the branch length effect: the higher the parsimony score for a branch, the longer the branch is likely to be in a probabilistic context, and the less its parsimony score should influence our preference among candidate trees. To justify the correction we use, it is helpful to consider the probability of one sequence evolving into another over some length of time *υ* under the JC69 model (Jukes and Cantor, 1969). This will be a product of two factors: the probability of the ending state being the same as the starting state, *p*_0_, and the probability of it being different, *p*_1_, both functions of *υ*. Say that we observe that *S* of *N* sites are different. Then the overall probability will be *p*_1_(*υ*)*^S^p*_0_(*υ*)*^N^*^−^*^S^* = (*p*_1_(*υ*)*/p*_0_(*υ*))*^S^p*_0_(*υ*)*^N^*. Thus, the overall probability is proportional to the ratio *p*_1_(*υ*)*/p*_0_(*υ*) raised to *S*, the number of sites that are different. The number of sites that are different is of course the same as the parsimony score. If *υ* is small, *p*_0_(*υ*) should be sufficiently close to 1 that we can ignore it. Assuming that the JC69 model is a reasonable approximation of more complex substitution models, we set 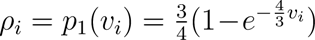, where *υ_i_* is the length of branch *i*. As an estimate of *υ_i_*, which can be computed easily, we use *υ̂_i_* = *S_i_/N* + *η*, where *N* is the total number of sites, and η is a small positive number (0.0001) to avoid *ρ_i_* = 0 when *S_i_* = 0. Table 1 gives some examples of pSPR1 and pSPR2 weights for different parsimony scores, branch lengths and choice of tuning parameters.

**Table 1:**
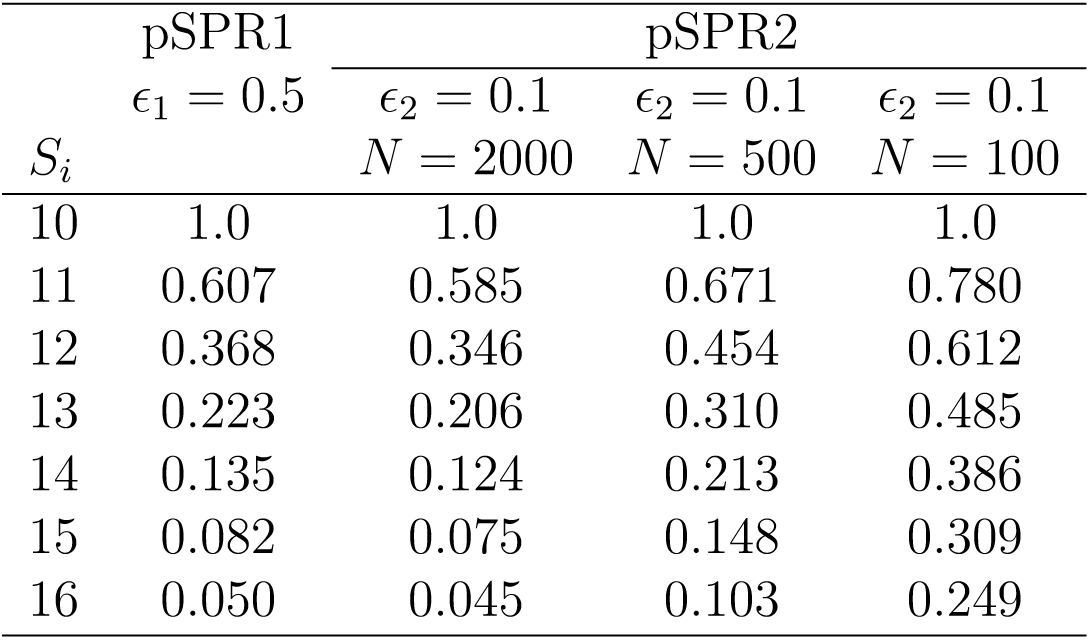
Examples of pSPR1 and pSPR2 weights (*ω_i_* values) for different branch lengths and parsimony scores given the tuning parameter values used here. All weights are scaled to the weight for a parsimony score of 10. Estimated branch lengths for pSPR2 are *υ̂_i_* = *S_i_/N* + 0.0001.

For all SPR moves, branch lengths are mapped from the old tree to the new tree as indicated by the branch labelling used above (Fig. 1). Then the lengths of the picked branch (*υ_a_*), the pendant branch moved with it (*υ_q_*), and the branch left behind (*υ_b_*), are each independently modified using a standard scaler move. These branch length changes modify the proposal ratio because the multiplier of the scaler move stretches parameter space, as explained in detail elsewhere (Holder et al., 2005).

### Tree bisection and reconnection (TBR)

The TBR moves pick an internal branch *a* at random, then prunes and regrafts each end of that branch in the corresponding subtree. The eTBR move applies the same extension mechanism as eSPR to both ends of *a*, guaranteeing that at least one randomly chosen end of the branch is moved from its original position (Lakner et al., 2008). The pTBR moves use the same weighting function as the pSPR moves (Equation 1), based on the parsimony score of the proposed new tree minus the sum of parsimony scores of the two subtrees resulting from bisecting the original tree at branch *a*. Using the same two choices for *ρ_i_* as in the pSPR case above results in two pTBR variants, pTBR1 and pTBR2.

For pTBR moves, evaluating all possible reconnection points would be computationally expensive in large trees. Therefore, we only considered candidate reconnection points maximally *δ* nodes away from *a*. For example, in Figure 1a, *c* or *p* is one node, and *d* or *r* is two nodes away from *a*. Thus, *δ* is an extra tuning parameter for the pTBR moves, besides *ϵ*.

For all TBR moves, branch lengths are mapped from the old tree to the new tree as for the SPR moves, and then the lengths of the chosen internal branch and the two pendant branches moved with it, one on each side, are modified using independent scaler moves in a fashion analogous to that of the SPR moves.

### Tuning Parameters

The tuning parameter values used for the tree moves in the analyses are summarized in Table 2. Choosing *p_e_* = 0.5 means that eSPR will propose NNI changes half the time (even more often if the extension mechanism is constrained by tips), and more radical changes the rest of the time. For eTBR, the same setting means that it will propose NNI changes 1*/*4 of the time. Choosing *δ* = 5 for a pTBR move generates up to approximately 2^6^2^6^ = 2^12^ = 4,096 candidate trees when there are no tip constraints. A pSPR move on a tree with *n* tips will generate up to roughly 2*n* candidate trees, that is, from around 700 to around 1,900 candidate trees for the empirical datasets analyzed here. The tuning parameter of the scaler moves used to modify branch lengths was set to 2 ln 1.05, yielding proposed changes of maximally 5% up or down in branch lengths (Larget and Simon, 1999).

**Table 2:**
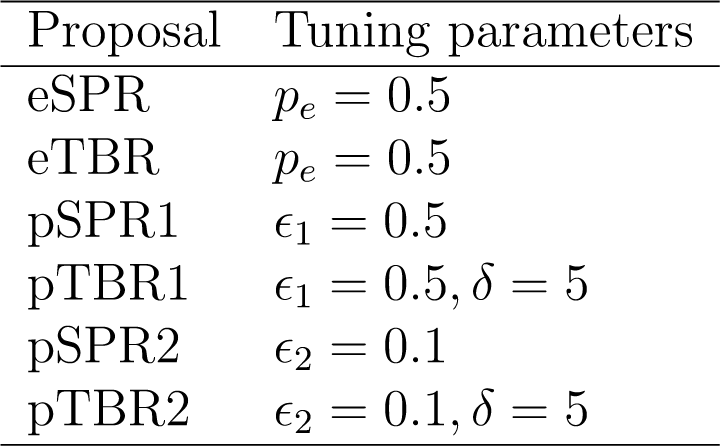
Tuning parameter values used here for the tree proposals. In addition to the tuning parameters listed in the table, all proposals used scaler moves with a tuning parameter of 2 ln 1.05 to change selected branch lengths.

### Simulations

We performed simulations to verify the implementation of the parsimony-guided tree proposals. Importantly, we tested whether the proposal ratio was correctly implemented so that we could retrieve the uniform prior distribution on topologies when the likelihood was set to be constant (zero).

The tree of five taxa used has two long branches separated by a short one (Fig. 2a). The parsimony score will favor the tree grouping the two long branches (C and D, Fig. 2b) when the sequence length approaches infinity, a phenomenon called long-branch attraction (LBA) (Felsenstein, 1978). We simulated sequences of 10,000 bp each for five taxa under the K80 model (Kimura, 1980) with *κ* = 4 on the true tree (Fig. 2a) using Seq-Gen 1.3.2x (Rambaut and Grassly, 1997). We generated ten simulated datasets, and verified that the tree inferred using maximum parsimony by PAUP* 4 beta 10 (Swofford, 2003) displayed LBA (Fig. 2b).

**Figure 2:**
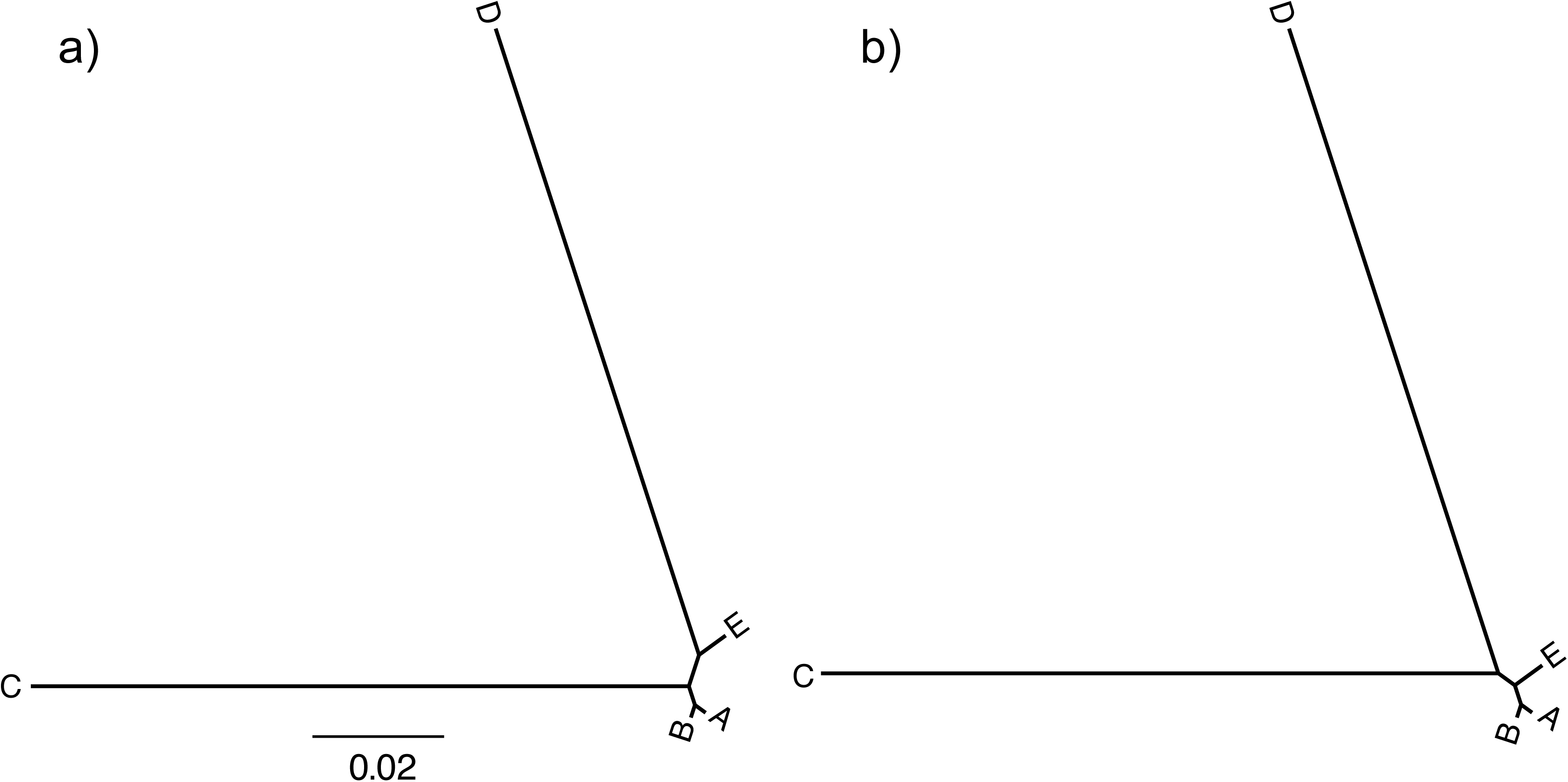
Simulation used to study the effect of long-branch attraction on the parsimony-guided tree proposals. (a) The true tree used in the simulation, with two long branches separated by a short one. (b) The tree inferred using maximum parsimony, erroneously joining long branches.

We used a uniform prior on topologies; thus, each topology had a prior probability of 1*/*15 ≈ 0.06667. For the parsimony-guided tree proposals, we used different numbers of sites in the sequence data (the first 0, 100, 1,000 or all 10,000 sites) to calculate the parsimony scores and the proposal ratio (Equation 2). With increasing number of sites, the parsimony weights will be more and more misleading in general, since no sequence data were used to compute the likelihood, but the proposal ratio should correct this imbalance.

In the MCMC runs, the only relevant model parameters are the topology and the branch lengths. We chose to combine each of the studied tree proposals with the default branch length proposal in MrBayes (a scaler move), using a 5:1 ratio of tree proposal to branch length proposal. For each combination, we ran a single chain for 10 million generations, without Metropolis coupling. The chain was sampled every 100 generations, and the first 25% of samples were discarded as burn-in.

### Empirical data

Evaluating the performance of tree proposals empirically is challenging. The problems need to be difficult enough to distinguish the performance of different topology proposals, while not being computationally too demanding to prohibit numerous repeated MCMC runs. Standard tree proposals under Metropolis-coupled MCMC are quite efficient in sampling from many small or simple tree spaces. Therefore, an obvious choice is to run empirical evaluations of tree proposals without Metropolis coupling, which makes the problem harder while bringing down the computational cost. The empirical datasets need to be reasonably large, with moderate to large number of taxa, as the parsimony-guided proposals should be particularly advantageous when sampling from large tree spaces. For this study, we chose to run the critical tests without Metropolis coupling but using reasonably large empirical datasets under a more realistic evolutionary model than JC69, namely the general time reversible model with gamma rate variation across sites (GTR + Γ_4_; Yang, 1994). We assumed a single partition to avoid any confounding influence of the partitioning scheme.

For empirical data, we do not know the true posterior distribution over trees. However, it may be possible to generate a reference sample of the tree space, which can be used as a reasonably accurate approximation of the true posterior. Here, we chose to start from a number of empirical datasets of suitable size. We then ran an analysis on each dataset for a long time using Metropolis coupling and a mix of tree proposals, in the hope of obtaining a sufficiently accurate approximation of the posterior, as indicated by the Average Standard Deviation of Split Frequencies (ASDSF) diagnostic (Lakner et al., 2008). We then used the datasets where we were able to obtain such accurate reference samples in studying the convergence and mixing of different tree proposals.

Specifically, we collected 20 datasets from TreeBase ranging from 344 to 934 taxa and from 1,740 to 16,542 sites (Table 3 and Supplementary Material, Table S1). Each dataset was treated as a single partition, and the evolutionary model was set to GTR+Γ_4_ (Yang, 1994). We used a uniform prior over tree topologies, and the prior for branch lengths was set to gamma-Dirichlet(1, 0.1, 1, 1) (Zhang et al., 2012). We used a flat Dirichlet prior for the exchangeability rates, fixed the stationary state frequencies to empirical, and used an Exponential(1.0) prior for the shape parameter of the discrete gamma distribution of rates across sites.

**Table 3:** Empirical datasets used in this study. The abbreviation (initial of first and last authors and publication year), number of taxa, number of sites, TreeBase study ID, and reference for each dataset are given. SQ10 was only found in GenBank (see Appendix of Savolainen et al., 2000).

| Abbr. | Taxa | Sites | ID | Reference |
| --- | --- | --- | --- | --- |
| SQ10 | 357 | 2925 | — | Savolainen et al. (2000) |
| DA10 | 357 | 4493 | S11008 | Davis and Anderson (2010) |
| LZ12 | 425 | 2468 | S12425 | Lu et al. (2012) |
| CL12 | 459 | 1740 | S12998 | Cardoso et al. (2012) |
| AZ12 | 545 | 5681 | S11973 | Aliscioni et al. (2012) |
| NP11 | 934 | 3675 | S11260 | Nagy et al. (2011) |

We first ran four independent runs with four Metropolis-coupled chains each (one cold and three heated) on each dataset using the default settings of MrBayes version 3.2.7. Specifically, we used the following mix of tree proposals: 1 sNNI : 2 eSPR : 1 eTBR : 2 pSPR2 : 1 pTBR2. The total probability of choosing a tree proposal from this mix was set to 46.67%, a branch length move to 50%, and a move changing a substitution model parameter to 3.33%. The details are available in the MrBayes scripts provided as Supplementary Material. We ran these analyses for 20 million generations each, sampling every 1,000 generations. The first 40% of samples were discarded as burn-in. The tree sample was considered as a good approximation of the posterior distribution when it had an ASDSF ≤ 0.02.

For the datasets where we could obtain convergence, we then ran 16 independent single-chain analyses (without Metropolis coupling) for each of three different mixes of the studied tree moves: (a) 2 eSPR : 1 eTBR; (b) 2 pSPR1 : 1 pTBR1; and (c) 2 pSPR2 : 1 pTBR2. In these runs, the probability of choosing a tree proposal from the mix was set to 36%, a branch length move to 60%, and a move changing a substitution model parameter to 4%. Each chain was run for 10 million generations starting from a random tree, and was sampled every 1,000 generations. The details are available in the MrBayes scripts provided as Supplementary Material.

To visualize the structure of the tree spaces, we used multidimensional scaling (MDS) based on the SPR distance between sampled trees (Whidden and Matsen, 2015). MDS methods compute a low-dimensional space (two-dimensional in our case) that represents a distance matrix (the minimum SPR distances between trees in our case) as accurately as possible. Calculating SPR distances among all sampled trees was prohibitively time consuming. Therefore, we randomly subsampled 4000 trees (∼10%) and calculated their pairwise SPR distances using the rspr software (Whidden et al., 2010, 2013). The sampled trees were then plotted in the resulting space.

## Results

### Simulations

When the likelihood was constant (no data), and no sites were used to compute the parsimony guide weights, all tree proposals tested were able to retrieve the uniform prior distribution over topologies with good accuracy (Fig. 3). When 100 sites (Fig. 4a) or 1,000 sites (Fig. 4b) were used to compute the parsimony weights, and likelihoods were artificially kept constant to test the ability of the proposal ratio to correct for the proposal bias, we still retrieved the uniform topology. However, when all 10,000 sites were included in computing the parsimony weights, some topologies were undersampled by the pSPR moves because of numerical inaccuracies in computing proposal ratios. Specifically, this affected the trees with the worst parsimony scores (topology 14 and 15 in Figure 4), the trees which are unlikely to be sampled at all in a real analysis because they also have low likelihood scores. Examining the simulated datasets revealed that the likelihood scores of topology 14 and 15 were around 300 log units lower than the scores of the best tree when all 10,000 sites were considered. This means that a correct MCMC algorithm sampling from the posterior should visit them considerably less than a googolth (10^−100^) as often as the best tree. Thus, for all practical purposes, the numerical errors affecting the sampling of these trees can be ignored. Importantly, the numerical errors did not cause oversampling of the LBA tree (topology 2) in relation to the best tree (topology 1) even under this extreme setting.

**Figure 3:**
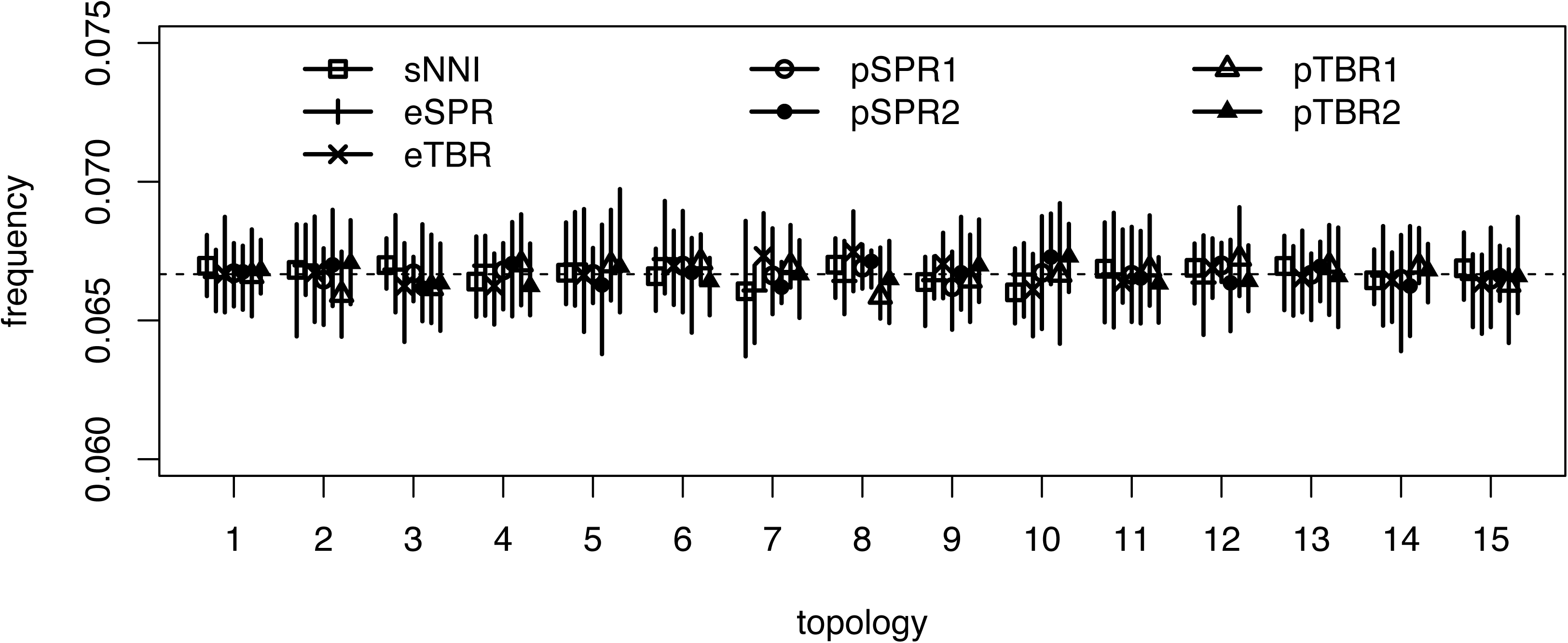
Test of the ability of the tree proposals to retrieve a uniform prior over topologies without data. We used a tree with five tips and a constant likelihood. The estimated posterior probabilities are represented as the mean (dot) and range (bar) across ten replicate analyses. The prior probability is 1/15 ≈ 0.06667 represented as horizontal dashed line. Topology 1 is the true tree (Fig. 2a) and topology 2 is the LBA tree (Fig. 2b).

**Figure 4:**
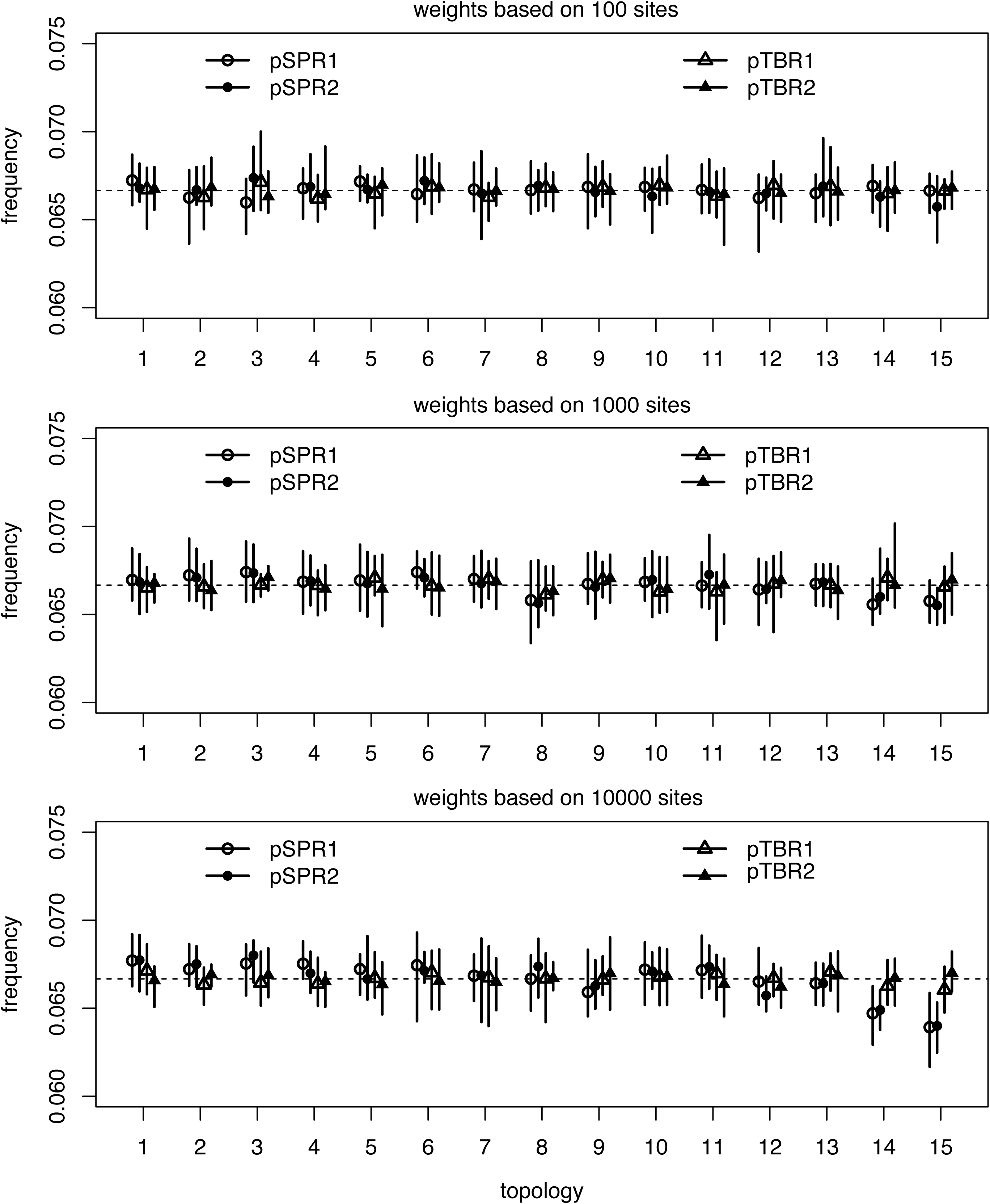
Test of the ability of the parsimony-guided tree proposals to retrieve a uniform prior over topologies when the first 100, 1000, or 10,000 sites were used to compute the parsimony weights. See legend to Figure 3.

Interestingly, the pTBR moves were not affected even when all 10,000 sites were included in computing parsimony weights (Fig. 4c). The reason is that they only pick interior branches, and therefore do not directly connect the worst trees (topology 14 and 15) with the best trees (topology 1 and 2) in a single move. This means that the pTBR proposal ratios are less extreme, making pTBR less sensitive to numerical errors in the computation of those ratios. However, this difference is of no consequence, as already the pSPR move is robust enough to numerical errors for all practical purposes.

### Empirical data

For six out of the 20 datasets, we were able to obtain reasonably accurate estimates of the posterior distribution in the reference runs (ASDSF ≤ 0.02, Table 4). Of the remaining 14 datasets, ASDSF values reached below 0.05 for three, and another three had ASDSF values below 0.10 (Supplementary Material, Table S2). For the rest of the datasets, the tree samples obtained in the four independent analyses were even more heterogeneous at 20 million generations, and it seemed for a few of them that the runs would have had to be extended considerably to obtain good estimates of the posterior distribution.

**Table 4:**
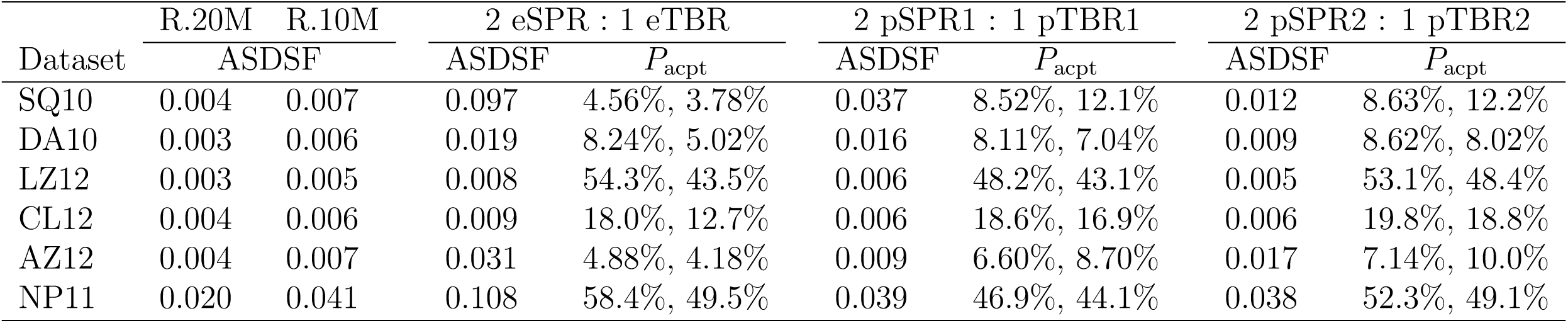
Results of reference runs (four runs with four Metropolis-coupled chains each) and test runs (16 single-chain runs) for each of the six datasets. We first give the average standard deviation of split frequencies (ASDSF) for the reference runs after 20 and 10 million generations (R.20M and R.10M, respectively). Then we give the ASDSF values and average acceptance proportion (*P*_acpt_) for 16 test runs using three different combinations of tree proposals. The reference tree samples after 20 million generations had ASDSF ≤ 0.02, and were considered as ground truth in the detailed studies of the convergence and mixing behavior of tree proposals.

Henceforth, we will focus on the six datasets for which we were able to generate reasonably accurate estimates of the posterior. Table 5 summarizes some characteristics of these datasets and the posteriors. The datasets span over a considerable range of tree shapes and percentages of missing data. Note that, for all six datasets, each tree topology in the posterior is sampled only once, so that the 95% credible sets all contain 45604 unique trees, that is, a unique tree for each sample from the posterior. This is because the posterior probability is spread out rather evenly over a very large number of topologies. Thus, our analyses only generate a limited subsample of the true credible set, which is likely to be much larger. Despite this shortcoming, the ASDSF diagnostic indicates that the reference runs have converged in the sense that the sampled trees are all representative of the true credible set in terms of split frequencies.

**Table 5:**
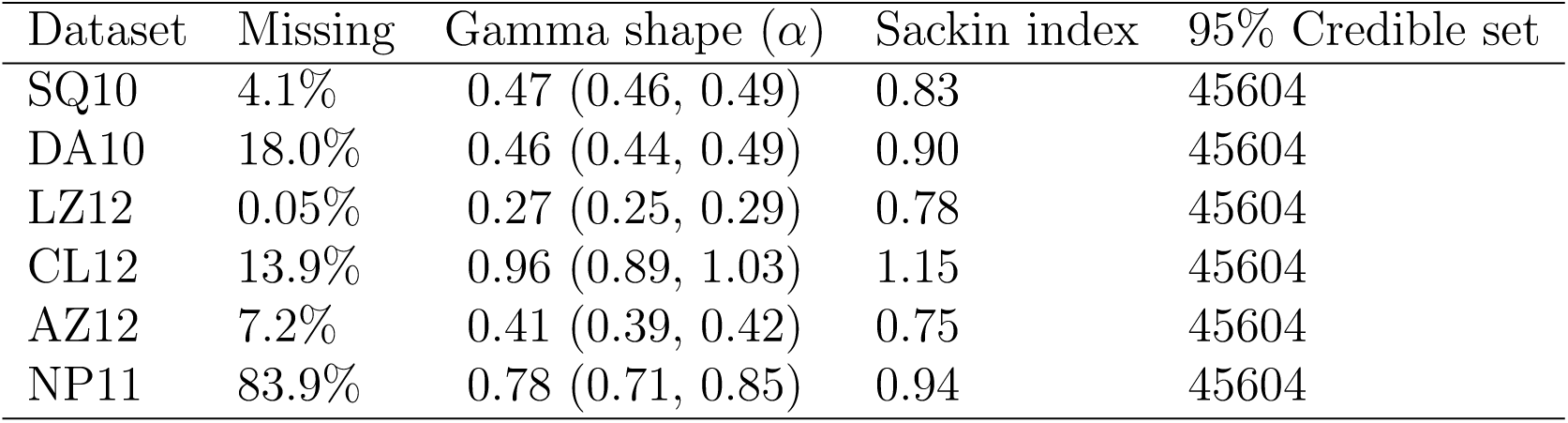
The percentage of missing data (including gaps), estimated gamma shape parameter (mean and 95% HPD interval), Sackin index (Mooers and Heard, 1997) measuring the balance of the consensus tree, and estimated size of the 95% credible set, for each of the six datasets. The Sackin index is normalized under the uniform model (Blum et al., 2006). Larger Sackin index indicates less balanced tree. The true credible sets are likely to be much larger; the run settings constrain the estimates to a maximum value. See text for further discussion.

We analyzed distances and computed tree spaces for a subsample of 4000 trees from each of the posterior samples. The minimum SPR distance among the trees in these subsamples ranged from 20 to over 200. The visualizations show that the tree spaces are quite different (Fig. 5). There are distinct islands of similar trees for datasets DA10 (two islands) and LZ12 (six islands grouped into two larger clusters). There is also an indication of island structure for SQ 10, while the posterior tree spaces are more homogeneous for the remaining three datasets (Fig. 5).

**Figure 5:**
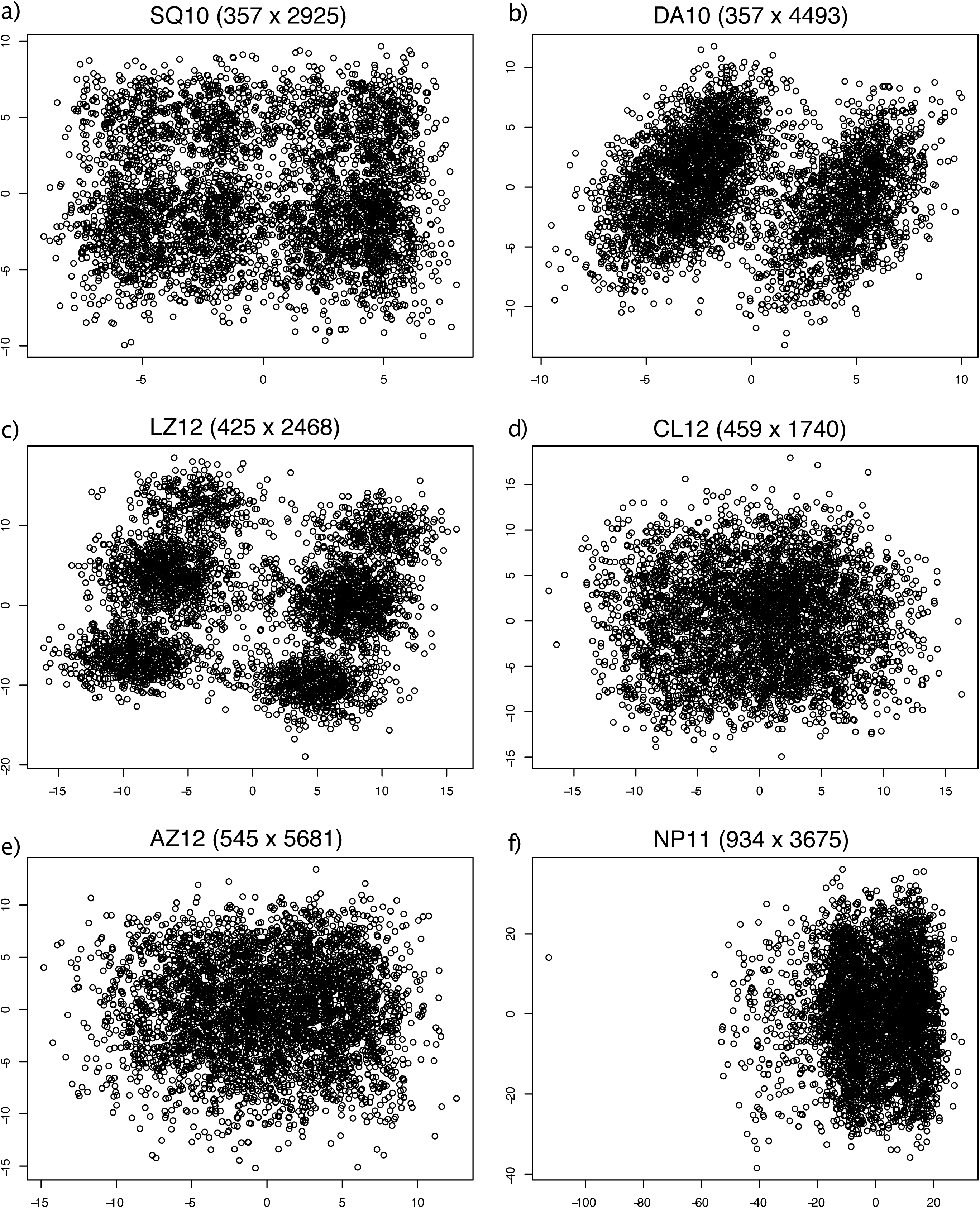
Multidimensional scaling to the SPR distance matrix of 4000 randomly subsampled trees from the posterior sample for each of the six datasets.

Note that these tree spaces are based on minimum SPR distances. Trees that appear close in this space may be more distant, or vice versa, in a space based on the actual probability of moving between them using, say, eSPR or pTBR. Nevertheless, the minimum SPR distances are presumably fairly similar to the true distances of the SPR and TBR moves considered here.

The single-chain test runs on the six datasets revealed striking differences in the performance of tree proposals. Analyses using parsimony-guided tree proposals (pSPR1+pTBR1 or pSPR2+pTBR2) increased in likelihood considerably faster than those using the standard extending proposals (eSPR+eTBR) (Fig. 6). The increase in likelihood was about an order of magnitude faster (note the logarithmic scale of the x-axis). There was no clear difference between the two variants of parsimony-guided proposals.

**Figure 6:**
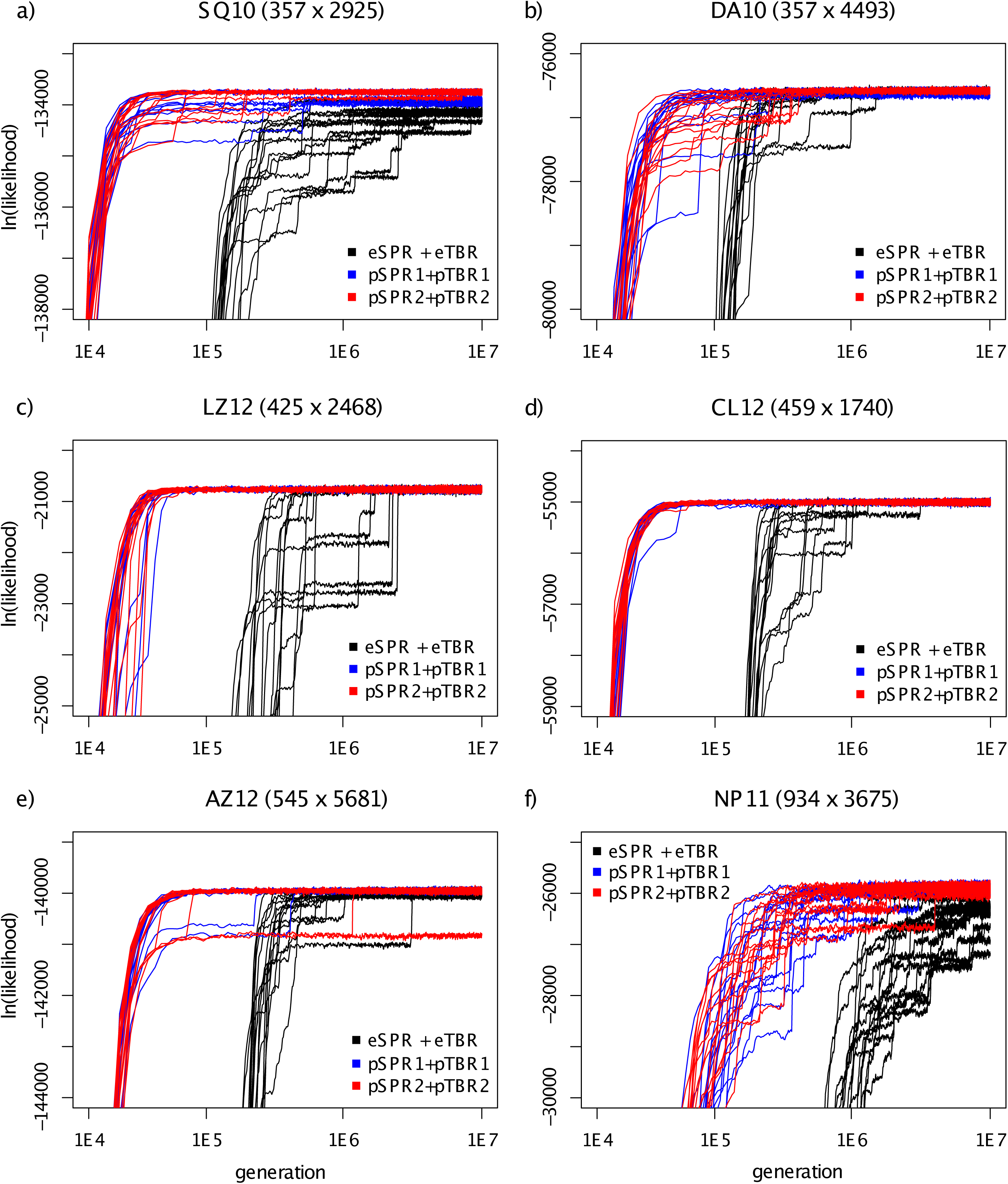
Performance of three different tree proposal combinations — eSPR+eTBR (black), pSPR1+pTBR1(blue) and pSPR2+pTBR2 (red) — on the six datasets (a–f) for which we had good reference samples from the posterior. We show the likelihood trace plots for 16 individual single-chain runs for each proposal combination. Note that all axes are in log scale.

Individual chains had a tendency to get stuck in suboptimal regions of tree space for a while before moving on towards more likely topologies, as indicated by local plateaus in the likelihood trace plots (Fig. 6). This phenomenon affected all tree proposals but was somewhat more pronounced in runs using standard moves, especially for datasets SQ10 and LZ12. For at least two of the datasets (SQ10 and NP11), the parsimony-guided chains reached higher likelihoods at the end of the test runs than all or most of the standard chains. The single exception was dataset AZ12, where one of the 32 parsimony-guided chains remained stuck throughout the run, despite the last of the 16 standard chains reaching the final plateau in log likelihoods one fourth into the run (Fig. 6e).

The convergence plots, which compare the topology samples from each of the test runs to the reference distribution using the ASDSF diagnostic, also reveal that the parsimony-guided proposals converged to the posterior much faster than the extending proposals, especially early on in the runs (Fig. 7; note the log scale on the y-axis). A few runs got stuck in local regions of tree space and did not cover the posterior distribution well; these runs generally corresponded to those that did not reach the final plateau in the likelihood trace plots (Fig. 6).

**Figure 7:**
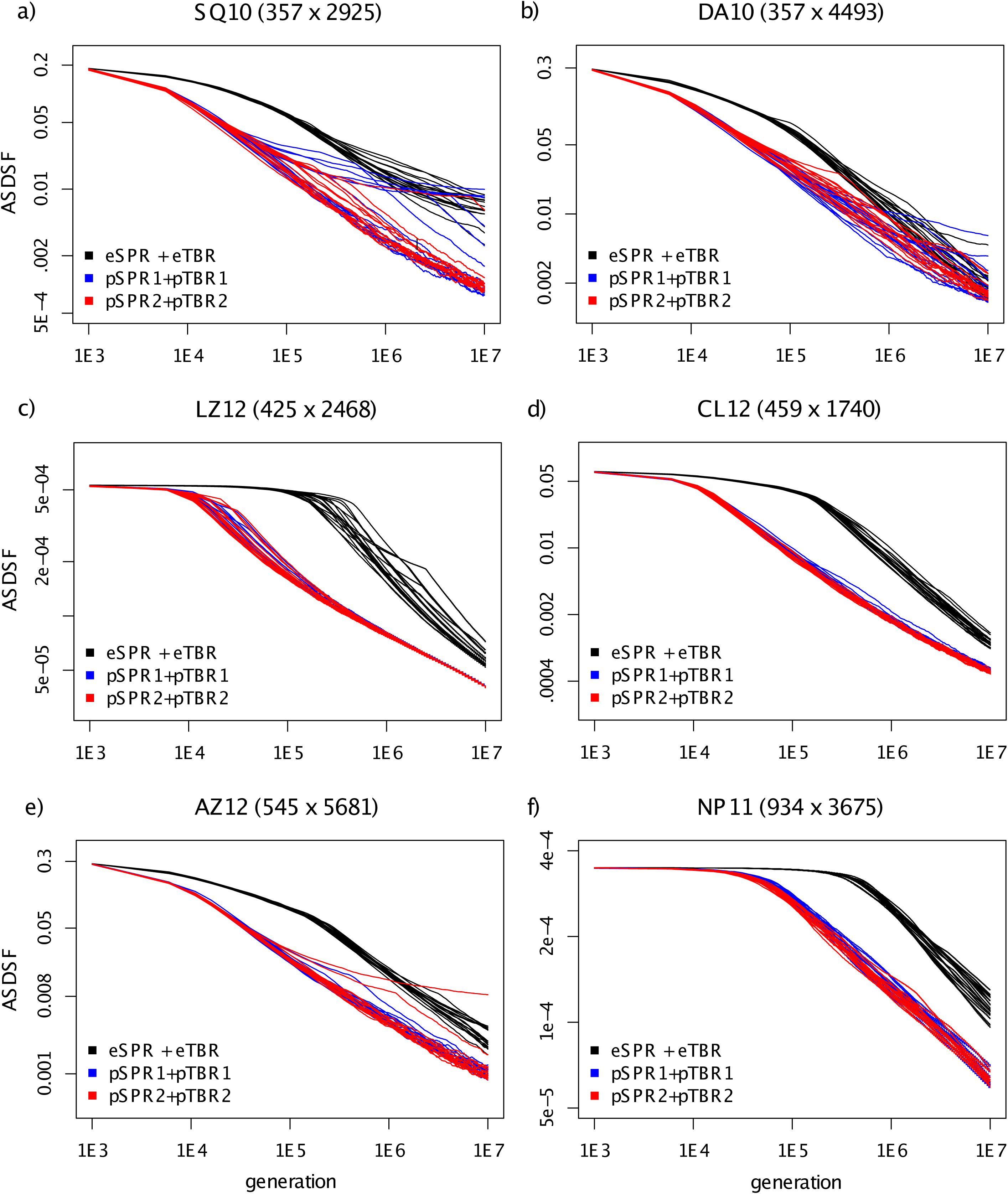
Performance of three different tree proposal combinations — eSPR+eTBR (black), pSPR1+pTBR1(blue) and pSPR2+pTBR2 (red) — on the six datasets (a–f) for which we had good reference samples from the posterior. We show the topological converge towards the reference sample as indicated by the ASDSF diagnostic for 16 individual single-chain runs for each proposal combination. Note that all axes are in log scale.

To look specifically at mixing behavior, we analyzed the last 5 million generations (the second half) of the runs. All of the chains had reached the same plateau in the likelihood plots at this point, except for the extending tree proposals on NP11, most of the chains on SQ10, and one parsimony-guided outlier on AZ12 (Fig. 8; note that the last 5 million generations correspond to only the last 10% of the x-axis in Fig. 7 because of the log scale). The mixing plots (Fig. 8) show that the parsimony-guided proposals usually covered the set of likely trees significantly faster than the extending proposals, even for samples drawn after the run was assumed to have converged to the posterior.

**Figure 8:**
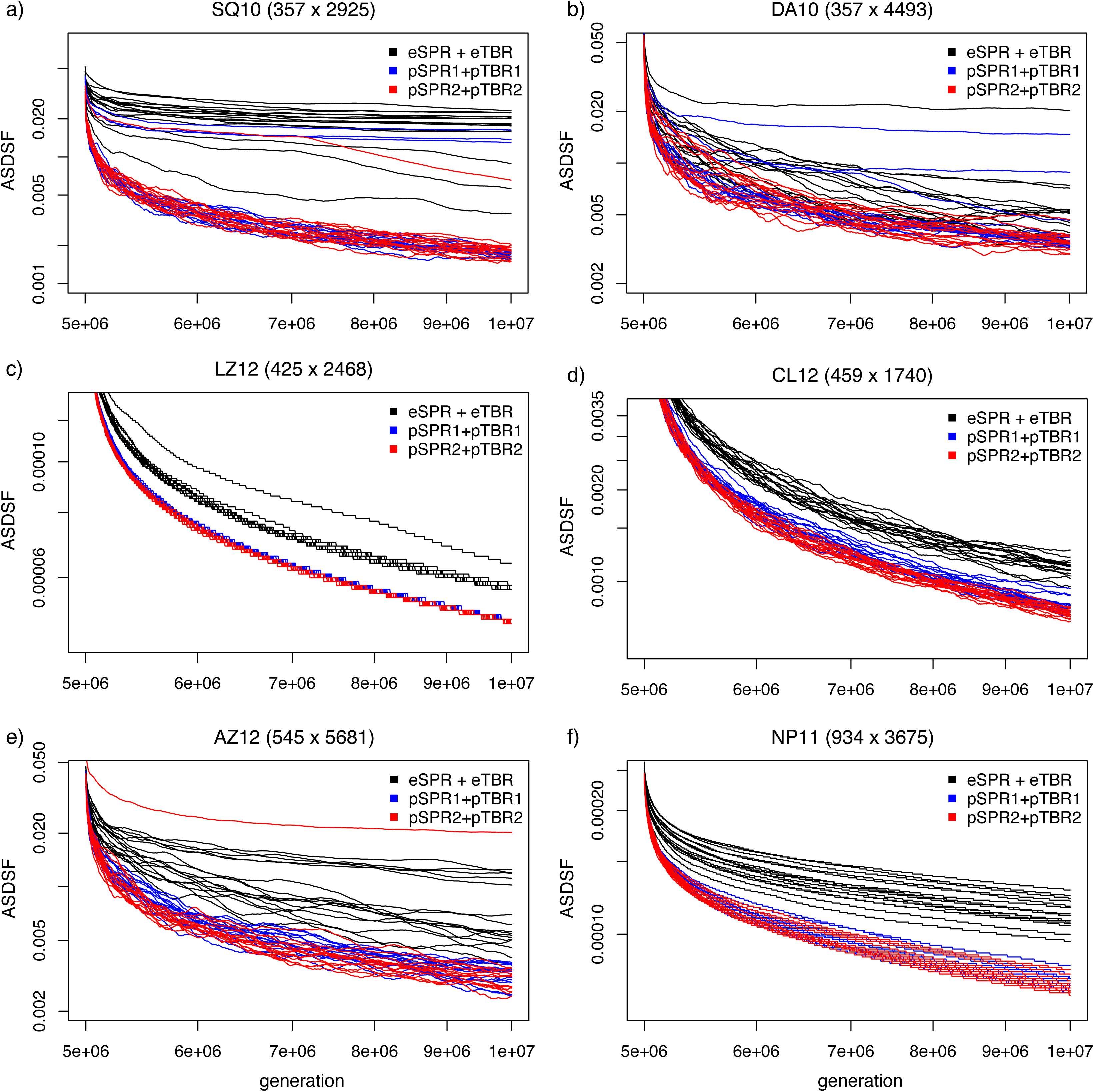
Mixing behavior of three different tree proposal combinations — eSPR+eTBR (black), pSPR1+pTBR1(blue) and pSPR2+pTBR2 (red) — on the six datasets (a–f) for which we had good reference samples from the posterior. We show the convergence towards the reference sample of topologies as indicated by the ASDSF diagnostic for 16 individual single-chain runs for each proposal combination. The plots correspond to the last 5 million generations of the runs shown in Figure 7, that is, the last 10% of the x-axis there. Thus, each run started with a tree with high posterior probability, and the rate at which the ASDSF drops represents the speed with which the chains cover the posterior. Note that the y-axis is in log scale but not the x-axis.

The mixing plots also reveal that tree spaces that were difficult to sample from using extending proposals (SQ10, DA10, AZ12), causing individual chains to get stuck for a very long time, were also difficult to sample from using parsimony-guided proposals. In general, however, it appeared that chains using parsimony-guided proposals were less likely to get stuck. The sampling difficulty is only partly correlated with the peakiness of the tree landscape indicated by the MDS plots (Fig. 5). Distinct islands in tree space can apparently cause chains to get stuck (DA10) but there are also tree spaces with distinct islands that are easy to sample from (LZ12) and spaces without obvious island structure that are difficult to sample from (AZ12).

## Discussion

Fast approximations of the posterior probability of different trees or tree topologies have the potential of making MCMC tree proposals “aware” of the structure of the local tree space around the current tree, allowing them to make smarter proposals with a higher chance of being accepted. The faster the approximation, the larger the region of tree space that a MCMC tree proposal could “see”, and the less likely it would get stuck on a local peak. Of course, speed typically comes at the expense of accuracy, so there is clearly a trade-off between these criteria when choosing an appropriate approximation method.

Parsimony scores are interesting in this context because they can be computed so rapidly. Evaluating the likelihood of an alignment of *c* discrete characters with *s* states over a tree with *n* tips has time complexity roughly proportional to *cns*^2^. Evaluating the parsimony score can be done in time proportional to *cn*, which is small in comparison. The constants involved in the time complexity equations are also considerably smaller for parsimony implementations, particularly with appropriate optimizations (Ronquist, 1998). Importantly, the parsimony scores of SPR and TBR candidate trees can be obtained in time that is dependent on only *c* for each candidate evaluated, once a *cn* order computation of parsimony sets has been completed. The parsimony sets need to be computed only once for the entire tree; after that they can be updated in time only dependent on *c*, under some reasonable assumptions about tree perturbations that are likely to be accepted in a hill-climbing or an MCMC algorithm (Ronquist, 1998). In this study, we used a naive implementation to compute parsimony scores, without employing any of the advanced optimizations described above. Nevertheless, we found that the pSPR and pTBR moves were only slightly slower than eSPR and eTBR. This means that there is considerable room for expansion of the region of tree space evaluated with parsimony scores beyond what we explored in the moves presented here.

The fact that parsimony scores may favor the wrong tree in some cases due to LBA has been widely discussed in the literature (Felsenstein, 1978). Therefore, one might be concerned that parsimony-guided proposals may introduce biases in the MCMC sample of trees. However, our simulations show that the parsimony-guided proposals are implemented correctly and that the proposal ratio adequately corrects for parsimony biases, such as the LBA effect. Even under extreme settings, when numerical inaccuracies start affecting the sampling of the least likely trees (the ones with the lowest parsimony weights but also with the lowest likelihood scores), we detect no oversampling of the LBA tree in relation to the best tree in our simulations.

Even though posterior distributions are sampled correctly, mismatches between the parsimony scores and posterior probabilities are likely to reduce the acceptance proportion and therefore the effectiveness of parsimony-guided proposals. It is possible that some of the variation in the success of parsimony-guided proposals that we observed is explained by differences in the effectiveness of the parsimony scores in approximating posterior probabilities for different datasets. However, such effects were not immediately apparent. One might expect datasets with more rate heterogeneity across sites to be more challenging than datasets with less variation in site rates, because rate heterogeneity would increase the mismatch between parsimony and likelihood scores. However, datasets that were the least challenging for parsimony-guided proposals included both the two datasets that had the least rate heterogeneity (CL12, NP11) and the dataset with the most pronounced rate variation across sites (LZ12; Table 5). One might also expect datasets with more variation in branch lengths to be more challenging for parsimony-guided proposals. However, there appeared to be no relation between variation in branch lengths (Fig. 9) and the difficulty of sampling the posterior correctly using parsimony-guided proposals. Of course, these conclusions are only based on six datasets, so they remain tentative at best.

**Figure 9:**
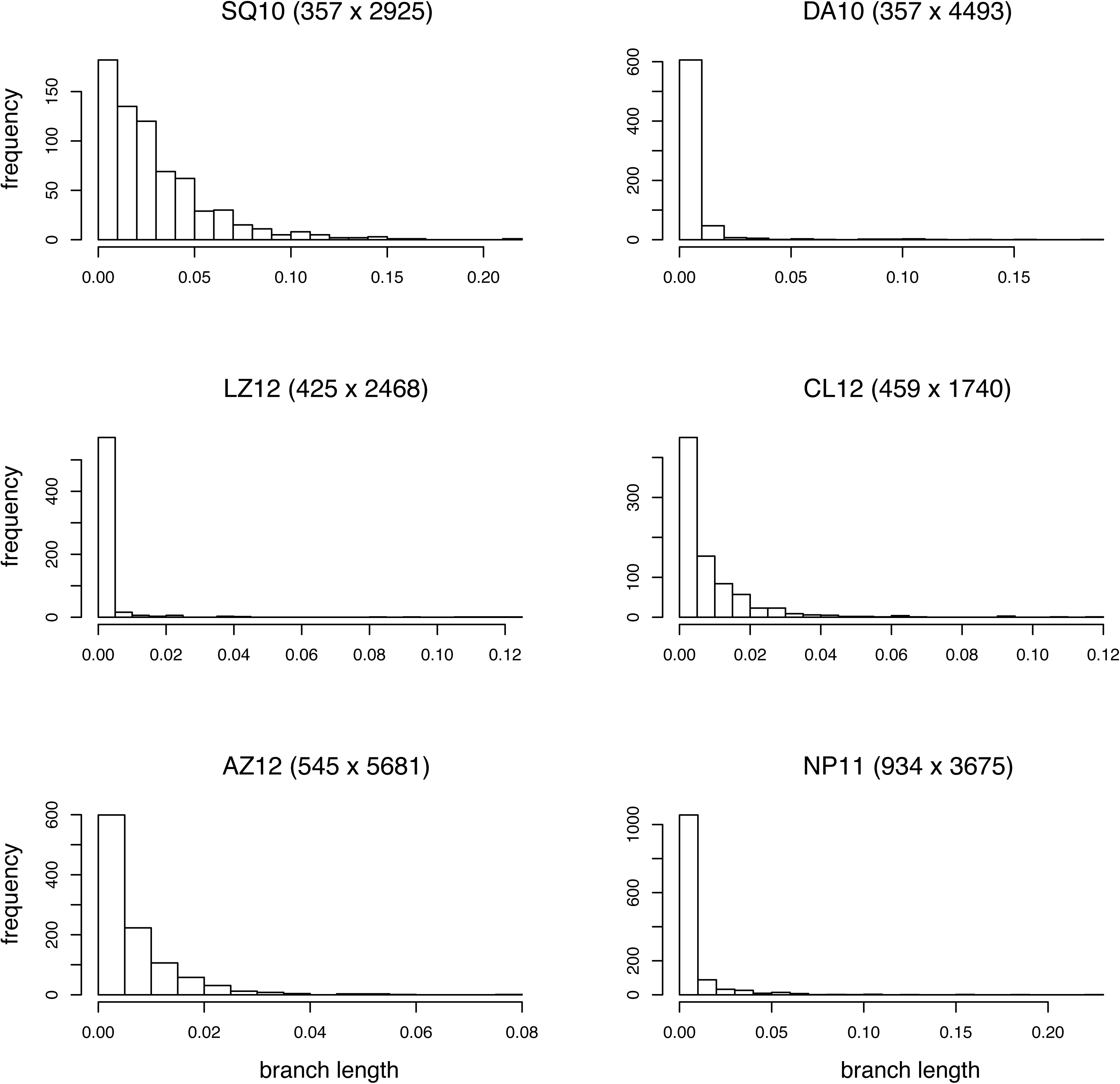
Histograms of the median branch lengths in the consensus tree of the reference tree samples for each of the six datasets.

In general, the tree spaces that were difficult to sample from using extending proposals were also comparatively more difficult to sample from using parsimony-guided proposals. Surprisingly, the peakiness of the tree space did not seem to be the only factor determining the sampling difficulty, as the more difficult datasets included both tree spaces with distinct tree islands (DA10; Fig. 5) and more homogeneous ones (AZ12). A potential explanation for this is that the tree space based on minimum SPR distances is different from the tree space based on the overall probability of moving between trees with high posterior probability using a particular tree proposal. It is the latter distance that actually determines the extent to which there are local peaks in the tree space for that proposal. Unfortunately, there are no methods that can visualize large tree spaces based on move probabilities as far as we know.

Intuitively, one would expect that parsimony-guided proposals should be particularly helpful in the burn-in phase of an MCMC run, when moving from a poor starting tree to a tree with high posterior probability. It seems less clear whether parsimony-guided proposals would also help chains mix well over the posterior once the best trees have been reached. Our results do show that parsimony-guided proposals can dramatically shorten the burn-in phase of an MCMC run, but they also show that parsimony-guided proposals accelerate mixing. Presumably, the ability of the parsimony-guided proposals to see promising peaks beyond the local region in tree space is important enough that it compensates for the mismatches between parsimony scores and posterior probabilities even during the later phases of an MCMC run.

As shown in Table 4, the ASDSF was always lower in the reference runs than in the test runs at 10 million generations, indicating that convergence and mixing were faster when using Metropolis coupling and combining the extending and parsimony-guided tree proposals. A question that arose during the study was whether this improvement in convergence was due largely to Metropolis coupling or to the combination of moves, To answer this question, we repeated some of the reference runs using the same mix of tree proposals, but without Metropolis coupling. The results (Supplementary Material, Table S3) indicate that the mix of proposals is responsible for a large proportion of the improvement in convergence. This supports the recommendation of using mixes of tree proposals in empirical analyses, which is also suggested by our observation that the relative performance of different tree proposals depends strongly on the dataset analyzed. These results also show that it is more computationally efficient to run long single runs than it is to use Metropolis coupling for these six datasets. This conclusion, of course, is not likely to hold for all datasets (see Whidden and Matsen, 2015).

The fact that we used a more realistic evolutionary model than the JC69 evolutionary model suggests that our results should be relevant for many empirical analyses, even though there is no guarantee that this is the case. Topological convergence may become either easier or harder for more realistic and complex evolutionary models, and the relative performance of different types of moves could change both with the model and the dataset analyzed.

Although we focused entirely on unrooted trees, our conclusions regarding the pSPR moves should apply also to rooted and clock-constrained trees. In fact, parsimony-guided moves have an interesting property that could be particularly useful when estimating divergence times in clock trees with fossils using total-evidence dating (also referred to as tip dating or integrative dating) (Ronquist et al., 2012a). In such analyses, fossil taxa tend to have an extremely large proportion of missing characters because they lack sequence data and usually have incomplete coding for morphology as well, while extant taxa can potentially be coded for all characters. A parsimony-guided proposal would then generate a much more diffuse proposal distribution when moving fossil taxa than when moving extant taxa, as would be appropriate.

Despite a large computational budget, our experiments were not able to distinguish clearly between the performance of the two different pSPR and pTBR variants. Clearly, there is a need for further experimentation with tuning parameters of the moves described here, and exploration of other types of approximations of the posterior tree probabilities. We also want to emphasize that, although our results were consistent across all the six datasets for which we were able to obtain reasonably accurate estimates of the posterior, the sample size is still small and it would be highly desirable to extend the experiments to more datasets. In either case, other experimental approaches than the one used here may be needed to make this practical. Although finding such approaches could prove challenging, our results indicate that efforts in this direction might be rewarding.

## Supplementary Material

Data available from the Dryad Data Repository: https://doi.org/10.5061/dryad.98mp657.

## Acknowledgments

This research was supported by Swedish Research Council (VR) Grant 2014-05901 (to F.R.). C.Z. is supported by the 100 Young Talents Program of Chinese Academy of Sciences and the Strategic Priority Research Program of Chinese Academy of Sciences (XDB26000000). Computations were performed using the Swedish National Infrastructure for Computing under projects SNIC 2014/1-323 and SNIC 2015/1-394.

## Appendix

### Metropolis-Hastings Tree Proposals

All tree proposals described in the paper are examples of the Metropolis-Hastings algorithm. It generates successive samples from the posterior distribution by proposing a new state in parameter space, *θ′*, conditional on the current state *θ*, according to a proposal distribution that we will denote *q*(*θ′* | *θ*). The new state is then accepted with a probability *R*, determined by the equation

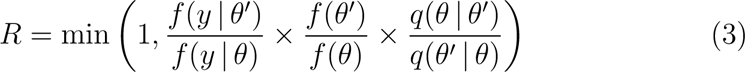

where the observations (data) are denoted *y*. The acceptance probability involves the product of three factors — the likelihood ratio (*f*(*y* |*θ′*)/*f*(*y* |*θ*)), the prior ratio (*f*(*θ′*)/*f*(*θ*)), and the proposal ratio (*q*(*θ* | *θ′*)/*q*(*θ′* | *θ*), also known as the Hastings ratio).

Note that the proposal ratio is computed as the probability of the backward move, returning to *θ* from *θ′*, divided by the probability of the forward move, proposing *θ′* when starting from *θ*. Assume we bias the move so that it is more probable to propose *θ′* from *θ* than the other way around. Then the proposal ratio for a move from *θ* to *θ*′ will be smaller than one, making it less likely to accept the move. However, if the likelihood of *θ′* is higher than that of *θ*, this preference can still pay off. A special case occurs if the preference for *θ′* over *θ* is exactly the same as the ratio of the posterior probabilities of these states (a Gibbs proposal). Then we have the Hastings ratio

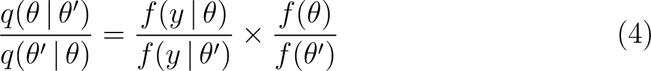

This is the reverse of the likelihood ratio times the prior ratio, which results in these terms cancelling the proposal ratio. Thus, the acceptance ratio becomes 1.0, which means that the proposal is always accepted, showing that a Gibbs proposal is a special case of the Metropolis-Hastings algorithm.

Computing the posterior probability ratio (the likelihood ratio times the prior ratio) for a set of candidate states is often computationally expensive. The goal we pursue in the current paper is to find proposals based on approximations of the posterior probability ratio that are easy to compute. For good approximations, the proposal ratio should come close to canceling the true posterior probability ratio, thereby increasing the average acceptance probability and improving mixing.

An unrooted phylogenetic tree *T* can be described as consisting of a topology *τ* and a set of branch lengths **v**. Typically, a phylogenetic model also includes substitution model parameters *σ*. Thus, we can write a state in parameter space as *θ* = (*τ*, **v**, *σ*). All tree proposals considered here leave *σ* unchanged, but propose a new topology *τ′* and (usually) a new set of branch lengths **v**′. The acceptance ratio is then obtained as

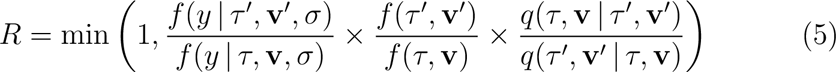

The computation of the likelihood ratio and the prior ratio in this equation is well known (e.g., see Felsenstein, 2003). Therefore, to describe a tree proposal, it is sufficient to specify how to generate a proposed state (*τ′*, **v**′) from the current state (*τ*, **v**) and how to compute the associated proposal ratio *q*(*τ*, **v** | *τ′*, **v**′)/*q*(*τ′*, **v**′ | *τ*, **v**).

### Implementation Details

For pSPR moves, calculating the proposal weight *ω_i_* (Equation 1) involves calculating the parsimony score *S_i_* of branch *i*. This score can be obtained easily by first computing the most parsimonious states at the root node of A (*z* in Fig. 1b) and at all nodes in subtree BCDE, then computing the number of most parsimonious changes among the three nodes incident to the potential regrafting point (*x*, *y* and *z* in Fig. 1b) using well-known algorithms (e.g., see Felsenstein, 2003). This procedure ensures that *S_i_* is the parsimony score of the tree after regrafting A at *i* minus the sum of the parsimony scores of the two subtrees A and BCDE. For pTBR moves, the *S_i_* scores are computed by first obtaining the most parsimonious states of all nodes in both subtrees, and then obtaining the number of most parsimonious changes among the four nodes, two in each subtree, adjacent to the proposed reconnection point.

Computationally, we do not obtain the proposal ratio (Equation 2) by directly calculating the *ω_i_* or their sum because this could result in severe numerical problems. Instead, to obtain the ratio associated with the backward or the forward move (the numerator or denominator in Equation 2), we first find the maximum value of *ω_i_* (assuming that 0 < *ρ_i_ <* 1) denoted as *ω_m_* = *ρ_m_^ϵS_m_^*. In pSPR1 or pTBR1, since *ρ_i_*’s are all equal to *e*^−1^, this is equivalent to finding the *ω* value for the minimal parsimony score. This may not be the case for pSPR2 or pTBR2, since the *ρ_i_*’s vary and *S_m_* may not be the minimal value among the *S_i_* values.

After finding *ω_m_*, we divide each *ω_i_* by *ω_m_* and convert *ω_i_*/*ω_m_* to

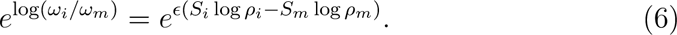

Using these transformed weights for the forward move and backward move allows us to obtain good numerical stability in computing the proposal ratio.

To further improve the precision of calculating the sum for *i* ∈ *B* \ *r* or *j* ∈ *B* \ *b*, we adopted the Kahan summation algorithm (Kahan, 1965). Even so, there might still be biases because of numerical inaccuracies in adding extremely unequal numbers. Take pSPR1 for example. Computing the proposal ratio for this move depends on weight terms of the form *e*^(*Sm*−*Si*)/2^ by default (*ϵ* = 0.5); see Equation 6. The maximum value of these terms is 1.0 when *S_m_* = *S_i_* (the most parsimonious tree) but the value can be extremely small when *S_m_* ≪ *S_i_* (e.g., *e*^−30^ when *S_m_* = 10 and *S_i_* = 70), making the small terms neglected during the summation (cf. Equation 2). This may affect moves considering both trees with extremely high and extremely low guide weights. The likely effect is undersampling of trees with extremely low guide weights, and distorted balance among trees with extremely low guide weights. In general, trees with extremely low guide weights also have extremely low likelihood scores. Therefore, the effect of these numerical errors should be negligible when analyzing empirical data.

## References

Aberer, A. J., K. Kobert, and A. Stamatakis. 2014. ExaBayes: massively parallel bayesian tree inference for the whole-genome era. Molecular Biology and Evolution 31:2553–2556.

Aliscioni, S., H. L. Bell, G. Besnard, P.-A. Christin, J. T. Columbus, M. R. Duvall, E. J. Edwards, L. Giussani, K. Hasenstab-Lehman, K. W. Hilu, T. R. Hodkinson, A. L. Ingram, E. A. Kellogg, S. Mashayekhi, O. Morrone, C. P. Osborne, N. Salamin, H. Schaefer, E. Spriggs, S. A. Smith, F. Zuloaga, and G. P. W. Grp II. 2012. New grass phylogeny resolves deep evolutionary relationships and discovers C4 origins. New Phytologist 193:304–312.

Blum, M. G. B., O. François, and S. Janson. 2006. The mean, variance and limiting distribution of two statistics sensitive to phylogenetic tree balance. The Annals of Applied Probability 16:2195–2214.

Bouchard-Côté, A., S. Sankararaman, and M. I. Jordan. 2012. Phylogenetic inference via sequential Monte Carlo. Systematic Biology 61:579–593.

Bouckaert, R., J. Heled, D. Kühnert, T. Vaughan, C.-H. Wu, D. Xie, M. A. Suchard, A. Rambaut, and A. J. Drummond. 2014. BEAST 2: a software platform for Bayesian evolutionary analysis. PLoS Computational Biology 10:e1003537.

Cardoso, D., L. P. de Queiroz, R. T. Pennington, H. C. de Lima, E. Fonty, M. F. Wojciechowski, and M. Lavin. 2012. Revisiting the Phylogeny of Papilionoid Legumes: New Insights From Comprehensively Sampled Early-Branching Lineages. American Journal of Botany 99:1991–2013.

Davis, C. C. and W. R. Anderson. 2010. A complete generic phylogeny of Malpighiaceae inferred from nucleotide sequence data and morphology. American Journal of Botany 97:2031–2048.

Drummond, A. J. and A. Rambaut. 2007. BEAST: Bayesian evolutionary analysis by sampling trees. BMC Evolutionary Biology 7:214.

Felsenstein, J. 1978. Cases in which parsimony or compatibility methods will be positively misleading. Systematic Biology 27:401–410.

Felsenstein, J. 2003. Inferring Phylogenies. Sinauer Associates, Sunderland, Massachusetts.

Geyer, C. J. 1991. Markov chain Monte Carlo maximum likelihood. Pp. 156–163 in E. M. Keramidas, ed. Computing Science and Statistics: Proc. 23rd Symp. Interface.

Hastings, W. K. 1970. Monte Carlo sampling methods using Markov chains and their applications. Biometrika 57:97–109.

Höhna, S., M. Defoin-Platel, and A. J. Drummond. 2008. Clock-constrained tree proposal operators in Bayesian phylogenetic inference. 8th IEEE International Conference on Bioinformatics and BioEngineering (BIBE) Pages 1–7.

Höhna, S. and A. J. Drummond. 2012. Guided tree topology proposals for Bayesian phylogenetic inference. Systematic Biology 61:1–11.

Höhna, S., M. J. Landis, T. A. Heath, B. Boussau, N. Lartillot, B. R. Moore, J. P. Huelsenbeck, and F. Ronquist. 2016. RevBayes: Bayesian phylogenetic inference using graphical models and an interactive model-specification language. Systematic Biology 65:726–736.

Holder, M. and P. O. Lewis. 2003. Phylogeny estimation: traditional and Bayesian approaches. Nature Reviews Genetics 4:275–284.

Holder, M. T., P. Lewis, D. Swofford, and B. Larget. 2005. Hastings ratio of the LOCAL proposal used in Bayesian phylogenetics. Systematic Biology 54:961–965.

Huelsenbeck, J. P., C. Ane, B. Larget, and F. Ronquist. 2008. A Bayesian perspective on a non-parsimonious parsimony model. Systematic Biology 57:406–419.

Huelsenbeck, J. P. and F. Ronquist. 2001. MRBAYES: Bayesian inference of phylogenetic trees. Bioinformatics 17:754–755.

Huelsenbeck, J. P., F. Ronquist, R. Nielsen, and J. P. Bollback. 2001. Bayesian inference of phylogeny and its impact on evolutionary biology. Science 294:2310–2314.

Jukes, T. H. and C. R. Cantor. 1969. Evolution of protein molecules. Mammalian Protein Metabolism Pages 21–132.

Kahan, W. 1965. Pracniques: further remarks on reducing truncation errors. Communications of the ACM 8:40.

Kimura, M. 1980. A simple method for estimating evolutionary rates of base substitutions through comparative studies of nucleotide sequences. Journal of Molecular Evolution 16:111–120.

Lakner, C., P. van der Mark, J. P. Huelsenbeck, B. Larget, and F. Ronquist. 2008. Efficiency of Markov chain Monte Carlo tree proposals in Bayesian phylogenetics. Systematic Biology 57:86–103.

Larget, B. 2013. The estimation of tree posterior probabilities using conditional clade probability distributions. Systematic Biology 62:501–511.

Larget, B. and D. Simon. 1999. Markov chain Monte Carlo algorithms for the Bayesian analysis of phylogenetic trees. Molecular Biology and Evolution 16:750–759.

Li, S., D. K. Pearl, and H. Doss. 2000. Phylogenetic tree construction using Markov chain Monte Carlo. Journal of the American Statistical Association 95:508.

Liu, J. S. 2004. Monte Carlo Strategies in Scientific Computing. Springer Series in Statistics Springer New York, New York.

Lu, B., Y. Zheng, R. W. Murphy, and X. Zeng. 2012. Coalescence patterns of endemic Tibetan species of stream salamanders (Hynobiidae: Batrachuperus). Molecular Ecology 21:3308–3324.

Mau, B. and M. A. Newton. 1997. Phylogenetic inference for binary data on dendrograms using Markov chain Monte Carlo. Journal of Computational and Graphical Statistics 6:122–131.

Metropolis, N., A. W. Rosenbluth, M. N. Rosenbluth, A. H. Teller, and E. Teller. 1953. Equation of state calculations by fast computing machines. Journal of Chemical Physics 21:1087–1092.

Mooers, A. O. and S. B. Heard. 1997. Inferring evolutionary process from phylogenetic tree shape. The Quarterly Review of Biology 72:31–54.

Nagy, L. G., T. Petkovits, G. M. Kovacs, K. Voigt, C. Vagvoelgyi, and T. Papp. 2011. Where is the unseen fungal diversity hidden? A study of Mortierella reveals a large contribution of reference collections to the identification of fungal environmental sequences. New Phytologist 191:789–794.

Nascimento, F. F., M. dos Reis, and Z. Yang. 2017. A biologist’s guide to Bayesian phylogenetic analysis. Nature Ecology & Evolution 1:1446–1454.

Peskun, P. H. 1973. Optimum Monte-Carlo sampling using Markov chains. Biometrika 60:607–612.

Rambaut, A. and N. C. Grassly. 1997. Seq-Gen: an application for the Monte Carlo simulation of DNA sequence evolution along phylogenetic trees. Computer Applications in the Biosciences: CABIOS 13:235–238.

Rannala, B. and Z. Yang. 1996. Probability distribution of molecular evolutionary trees: a new method of phylogenetic inference. Journal of Molecular Evolution 43:304–311.

Ronquist, F. 1998. Fast Fitch-Parsimony Algorithms for Large Data Sets. Cladistics 14:387–400.

Ronquist, F. and J. P. Huelsenbeck. 2003. MrBayes 3: Bayesian phylogenetic inference under mixed models. Bioinformatics 19:1572–1574.

Ronquist, F., J. P. Huelsenbeck, and T. Britton. 2004. Bayesian Supertrees. Pages 193–224 in Phylogenetic Supertrees. Springer, Dordrecht, Dordrecht.

Ronquist, F., S. Klopfstein, L. Vilhelmsen, S. Schulmeister, D. L. Murray, and A. P. Rasnitsyn. 2012a. A total-evidence approach to dating with fossils, applied to the early radiation of the Hymenoptera. Systematic Biology 61:973–999.

Ronquist, F., M. Teslenko, P. van der Mark, D. L. Ayres, A. Darling, S. Höhna, B. Larget, L. Liu, M. A. Suchard, and J. P. Huelsenbeck. 2012b. MrBayes 3.2: efficient Bayesian phylogenetic inference and model choice across a large model space. Systematic Biology 61:539–542.

Savolainen, V., M. W. Chase, S. B. Hoot, C. M. Morton, D. E. Soltis, C. Bayer, M. F. Fay, A. Y. de Bruijn, S. Sullivan, and Y. L. Qiu. 2000. Phylogenetics of flowering plants based on combined analysis of plastid atpB and rbcL gene sequences. Systematic Biology 49:306–362.

Swofford, D. L. 2003. PAUP*. Phylogenetic Analysis Using Parsimony (*and Other Methods). Version 4. Sinauer Associates, Sunderland, Massachusetts.

Wang, L., A. Bouchard-Côté, and A. Doucet. 2016. Bayesian Phylogenetic Inference Using a Combinatorial Sequential Monte Carlo Method. Journal of the American Statistical Association 110:1362–1374.

Whidden, C., R. G. Beiko, and N. Zeh. 2010. Fast FPT Algorithms for Computing Rooted Agreement Forests: Theory and Experiments. Pages 141–153 in Proceedings of the 9th International Conference on Experimental Algorithms SEA’10 Springer-Verlag, Berlin, Heidelberg.

Whidden, C., R. G. Beiko, and N. Zeh. 2013. Fixed-Parameter Algorithms for Maximum Agreement Forests. SIAM Journal on Computing 42:1431–1466.

Whidden, C. and F. A. Matsen. 2015. Quantifying MCMC exploration of phylogenetic tree space. Systematic Biology 64:472–491.

Yang, Z. 1994. Maximum likelihood phylogenetic estimation from DNA sequences with variable rates over sites: approximate methods. Journal of Molecular Evolution 39:306–314.

Yang, Z. 2014. Molecular Evolution: A Statistical Approach. Oxford University Press, Oxford, UK.

Yang, Z. and B. Rannala. 1997. Bayesian phylogenetic inference using DNA sequences: a Markov Chain Monte Carlo Method. Molecular Biology and Evolution 14:717–724.

Yang, Z. and B. Rannala. 2012. Molecular phylogenetics: principles and practice. Nature Reviews Genetics 13:303–314.

Zhang, C., B. Rannala, and Z. Yang. 2012. Robustness of compound Dirichlet priors for Bayesian inference of branch lengths. Systematic Biology 61:779–784.

