## Supplementary Material for "Using Parsimony-Guided Tree Proposals to Accelerate Convergence in Bayesian Phylogenetic Inference"

December 28, 2018

Running Head: PARSIMONY-GUIDED TREE PROPOSALS

[1]*Key Laboratory of Vertebrate Evolution and Human Origins, Institute of Vertebrate Paleontology and Paleoanthropology, Chinese Academy of Sciences, Beijing 100044, China;*
[2]*Center for Excellence in Life and Paleoenvironment, Chinese Academy of Sciences, Beijing 100044, China;*
[3]*Department of Integrative Biology, University of California, Berkeley, CA 94720, USA;*
[4]*Department of Bioinformatics and Genetics, Swedish Museum of Natural History, Box 50007, SE-10405 Stockholm, Sweden;*

### Supplementary Material

#### MrBayes scripts used in the analyses

```
Begin mrbayes;
  set user=devel;

  [reference run]
  exe /path/to/mydata.nex;
  lset nst=6 rates=gamma;
  prset statefreqpr=fixed(empirical);
  mcmcp nrun=4 nchain=4 ngen=20000000 samplefreq=1000;
  propset  ParsSPR(Tau,V)$prob=0;
  propset      NNI(Tau,V)$prob=4;
  propset   ExtSPR(Tau,V)$prob=8;
  propset   ExtTBR(Tau,V)$prob=4;
  propset ParsSPR2(Tau,V)$prob=8;
  propset ParsTBR2(Tau,V)$prob=4;
  mcmc filename=mydata.ref;
  sumt burninfrac=0.4;

  [test run 1: eSPR+eTBR]
  exe /path/to/mydata.nex;
  lset nst=6 rates=gamma;
  prset statefreqpr=fixed(empirical);
  mcmcp nrun=16 nchain=1 ngen=10000000 samplefreq=1000;
  propset  ParsSPR(Tau,V)$prob=0;
  propset      NNI(Tau,V)$prob=0;
  propset   ExtSPR(Tau,V)$prob=12;
  propset   ExtTBR(Tau,V)$prob=6;
  mcmc filename=mydata.test1;

  [test run 2: pSPR1+pTBR1]
  exe /path/to/mydata.nex;
  lset nst=6 rates=gamma;
  prset statefreqpr=fixed(empirical);
  mcmcp nrun=16 nchain=1 ngen=10000000 samplefreq=1000;
  propset  ParsSPR(Tau,V)$prob=0;
```

```
  propset      NNI(Tau,V)$prob=0;
  propset   ExtSPR(Tau,V)$prob=0;
  propset   ExtTBR(Tau,V)$prob=0;
  propset ParsSPR1(Tau,V)$prob=12;
  propset ParsTBR1(Tau,V)$prob=6;
  mcmc filename=mydata.test2;

  [test run 3: pSPR2+pTBR2]
  exe /path/to/mydata.nex;
  lset nst=6 rates=gamma;
  prset statefreqpr=fixed(empirical);
  mcmcp nrun=16 nchain=1 ngen=10000000 samplefreq=1000;
  propset  ParsSPR(Tau,V)$prob=0;
  propset      NNI(Tau,V)$prob=0;
  propset   ExtSPR(Tau,V)$prob=0;
  propset   ExtTBR(Tau,V)$prob=0;
  propset ParsSPR2(Tau,V)$prob=12;
  propset ParsTBR2(Tau,V)$prob=6;
  mcmc filename=mydata.test3;

  [compare a test with the reference]
  compareref filename1=mydata.test1.run1.t filename2=mydata.ref
    outputname=compareref.test1.run1.txt;
  compareref filename1=mydata.test1.run2.t filename2=mydata.ref
    outputname=compareref.test1.run2.txt;
  ...
  compareref filename1=mydata.test3.run16.t filename2=mydata.ref
    outputname=compareref.test3.run16.txt;
End;
```

Table S1: Empirical datasets selected in preliminary tests but not used in the main study. The abbreviation (initial of first and last authors and publication year), number of taxa, number of sites, TreeBase study ID, and reference for each dataset are given.

| Abbr. | Taxa | Sites | ID | Reference |
|---|---|---|---|---|
| SW09 | 344 | 9800 | S12224 | Svenson and Whiting (2009) |
| SG13 | 400 | 6299 | S11736 | Schoch and Grube (2013) |
| EC10 | 413 | 3632 | S10454 | Evans et al. (2010) |
| SS09 | 434 | 7062 | S2137 | Schoch et al. (2009) |
| QS13 | 445 | 2336 | S14373 | Qi et al. (2013) |
| LS14 | 477 | 5167 | S15950 | Laenen et al. (2014) |
| KN11 | 492 | 9398 | S11336 | Katz et al. (2011) |
| MM14 | 510 | 4044 | S13659 | Mishler et al. (2014) |
| MM13 | 620 | 2273 | S13670 | McFrederick et al. (2013) |
| HS14 | 684 | 6537 | S16248 | Huvet and Stumpf (2014) |
| FS11 | 699 | 6950 | S11116 | Fiz-Palacios et al. (2011) |
| HP12 | 736 | 16542 | S13572 | Hermant et al. (2012) |
| PW11 | 767 | 5814 | S10982 | Pyron et al. (2011) |
| SL14 | 858 | 9960 | S15260 | Siu-Ting et al. (2014) |

Table S2: Results of reference runs (four runs with four Metropolis-coupled chains each) and test runs (16 single-chain runs) for each of the 14 datasets. We first give the average standard deviation of split frequencies (ASDSF) for the reference runs after 20 and 10 million generations (R.20M and R.10M, respectively). Then we give the ASDSF values and average acceptance proportion ($P_{\text{acpt}}$) for 16 test runs using three different combinations of tree proposals. The reference tree samples all have ASDSF ¿ 0.02, thus were not used in detailed studies of the convergence and mixing behavior of tree proposals.

| | R.20M | R.10M | 2 eSPR : 1 eTBR | | 2 pSPR1 : 1 pTBR1 | | 2 pSPR2 : 1 pTBR2 | |
|---|---|---|---|---|---|---|---|---|
| DataSet | ASDSF | | ASDSF | $P_{\text{acpt}}$ | ASDSF | $P_{\text{acpt}}$ | ASDSF | $P_{\text{acpt}}$ |
| SW09 | 0.089 | 0.102 | 0.162 | 0.63%, 0.55% | 0.200 | 0.53%, 0.73% | 0.211 | 0.55%, 0.78% |
| SG13 | 0.035 | 0.046 | 0.163 | 3.24%, 2.33% | 0.245 | 1.76%, 1.39% | 0.235 | 1.83%, 1.49% |
| EC10 | 0.129 | 0.136 | 0.218 | 3.65%, 2.48% | 0.186 | 3.92%, 3.20% | 0.176 | 4.15%, 3.38% |
| SS09 | 0.038 | 0.055 | 0.090 | 13.2%, 7.63% | 0.073 | 5.68%, 2.80% | 0.090 | 5.65%, 2.78% |
| QS13 | 0.122 | 0.150 | 0.205 | 6.49%, 4.09% | 0.208 | 3.54%, 2.01% | 0.223 | 3.59%, 2.08% |
| LS14 | 0.171 | 0.186 | 0.279 | 7.42%, 5.31% | 0.256 | 8.74%, 7.33% | 0.228 | 9.79%, 8.94% |
| KN11 | 0.167 | 0.206 | 0.236 | 3.21%, 2.48% | 0.234 | 3.26%, 2.53% | 0.231 | 3.25%, 2.43% |
| MM14 | 0.063 | 0.070 | 0.189 | 10.9%, 7.62% | 0.134 | 6.54%, 4.56% | 0.126 | 7.18%, 5.07% |
| MM13 | 0.185 | 0.186 | 0.232 | 15.4%, 11.4% | 0.204 | 13.0%, 10.0% | 0.217 | 13.5%, 10.7% |
| HS14 | 0.044 | 0.056 | 0.152 | 9.01%, 5.76% | 0.083 | 10.2%, 8.93% | 0.088 | 11.0%, 9.79% |
| FS11 | 0.151 | 0.166 | 0.232 | 5.37%, 3.71% | 0.207 | 2.93%, 3.29% | 0.203 | 2.98%, 3.43% |
| HP12 | 0.118 | 0.118 | 0.107 | 36.3%, 27.6% | 0.146 | 20.3%, 9.76% | 0.147 | 20.4%, 9.83% |
| PW11 | 0.080 | 0.104 | 0.171 | 4.59%, 3.33% | 0.195 | 2.49%, 2.28% | 0.190 | 2.49%, 2.34% |
| SL14 | 0.140 | 0.154 | 0.206 | 5.19%, 3.62% | 0.228 | 1.75%, 1.27% | 0.227 | 1.57%, 1.23% |

Table S3: Topological convergence (ASDSF values) of the reference analysis using Metropolis coupling ($(MC)^3$) compared to that of an equivalent set of runs using the same mix of proposals but without Metropolis coupling (MCMC) at 10 and 20 million generations. The datasets analyzed here are the ones for which the reference analyses generated good samples of the posterior distribution (ASDSF $\leq 0.02$).

| | Reference $(MC)^3$ | | MCMC | |
|---|---|---|---|---|
| Dataset | 20M | 10M | 20M | 10M |
| SQ10 | 0.004 | 0.007 | 0.005 | 0.007 |
| DA10 | 0.003 | 0.006 | 0.006 | 0.008 |
| LZ12 | 0.003 | 0.005 | 0.004 | 0.006 |
| CL12 | 0.004 | 0.006 | 0.004 | 0.006 |
| AZ12 | 0.004 | 0.007 | 0.006 | 0.007 |
| NP11 | 0.020 | 0.041 | 0.027 | 0.038 |

### References


Evans, N. M., M. T. Holder, M. S. Barbeitos, B. Okamura, and P. Cartwright. 2010. The Phylogenetic Position of Myxozoa: Exploring Conflicting Signals in Phylogenomic and Ribosomal Data Sets. Molecular Biology and Evolution 27:2733–2746.

Fiz-Palacios, O., H. Schneider, J. Heinrichs, and V. Savolainen. 2011. Diversification of land plants: insights from a family-level phylogenetic analysis. BMC Evolutionary Biology 11:341.

Hermant, M., F. Hennion, I. V. Bartish, B. Yguel, and A. Prinzing. 2012. Disparate relatives: Life histories vary more in genera occupying intermediate environments. Perspectives in Plant Ecology Evolution and Systematics 14:283–301.

Huvet, M. and M. P. H. Stumpf. 2014. Overlapping genes: a window on gene evolvability. BMC Genomics 15:721.

Katz, L. A., J. Grant, L. W. Parfrey, A. Gant, C. J. O'Kelly, O. R. Anderson, R. E. Molestina, and T. Nerad. 2011. Subulatomonas tetraspora nov gen. nov sp is a Member of a Previously Unrecognized Major Clade of Eukaryotes. Protist 162:762–773.

Laenen, B., B. Shaw, H. Schneider, B. Goffinet, E. Paradis, A. Desamore, J. Heinrichs, J. C. Villarreal, S. R. Gradstein, S. F. McDaniel, D. G. Long, L. L. Forrest, M. L. Hollingsworth, B. Crandall-Stotler, E. C. Davis, J. Engel, M. Von Konrat, E. D. Cooper, J. Patino, C. J. Cox, A. Vanderpoorten, and A. J. Shaw. 2014. Extant diversity of bryophytes emerged from successive post-Mesozoic diversification bursts. Nature Communications 5.

McFrederick, Q. S., J. J. Cannone, R. R. Gutell, K. Kellner, R. M. Plowes, and U. G. Mueller. 2013. Specificity between Lactobacilli and Hymenopteran Hosts Is the Exception Rather than the Rule. Applied and Environmental Microbiology 79:1803–1812.

Mishler, B. D., N. Knerr, C. E. Gonzalez-Orozco, A. H. Thornhill, S. W. Laffan, and J. T. Miller. 2014. Phylogenetic measures of biodiversity and neo- and paleo-endemism in Australian Acacia. Nature Communications 5:4473.

Pyron, R. A., F. T. Burbrink, G. R. Colli, A. N. Montes de Oca, L. J. Vitt, C. A. Kuczynski, and J. J. Wiens. 2011. The phylogeny of advanced snakes (Colubroidea), with discovery of a new subfamily and comparison of support methods for likelihood trees. Molecular Phylogenetics and Evolution 58:329–342.

Qi, X., A. S. Chanderbali, G. K.-S. Wong, D. E. Soltis, and P. S. Soltis. 2013. Phylogeny and evolutionary history of glycogen synthase kinase 3/SHAGGY-like kinase genes in land plants. BMC Evolutionary Biology 13:143.

Schoch, C. L. and M. Grube. 2013. Pezizomycotina: Dothideomycetes and Arthoniomycetes. Systematics and Evolution: Part B (The Mycota) .

Schoch, C. L., G.-H. Sung, F. López-Giráldez, J. P. Townsend, J. Miadlikowska, V. Hofstetter, B. Robbertse, P. B. Matheny, F. Kauff, Z. Wang, C. Gueidan, R. M. Andrie, K. Trippe, L. M. Ciufetti, A. Wynns, E. Fraker, B. P. Hodkinson, G. Bonito, J. Z. Groenewald, M. Arzanlou, G. S. de Hoog, P. W. Crous, D. Hewitt, D. H. Pfister, K. Peterson, M. Gryzenhout, M. J. Wingfield, A. Aptroot, S.-O. Suh, M. Blackwell, D. M. Hillis, G. W. Griffith, L. A. Castlebury, A. Y. Rossman, H. T. Lumbsch, R. Lücking, B. Büdel, A. Rauhut, P. Diederich, D. Ertz, D. M. Geiser, K. Hosaka, P. Inderbitzin, J. Kohlmeyer, B. Volkmann-Kohlmeyer, L. Mostert, K. O'Donnell, H. Sipman, J. D. Rogers, R. A. Shoemaker, J. Sugiyama, R. C. Summerbell, W. Untereiner, P. R. Johnston, S. Stenroos, A. Zuccaro, P. S. Dyer, P. D. Crittenden, M. S. Cole, K. Hansen, J. M. Trappe, R. Yahr, F. Lutzoni, and J. W. Spatafora. 2009. The Ascomycota Tree of Life: A Phylum-wide Phylogeny Clarifies the Origin and Evolution of Fundamental Reproductive and Ecological Traits. Systematic Biology 58:224–239.

Siu-Ting, K., D. J. Gower, D. Pisani, R. Kassahun, F. Gebresenbet, M. Menegon, A. A. Mengistu, S. A. Saber, R. de Sa, M. Wilkinson, and S. P. Loader. 2014. Evolutionary relationships of the Critically Endangered frog Ericabatrachus baleensis Largen, 1991 with notes on incorporating previously unsampled taxa into large-scale phylogenetic analyses. BMC Evolutionary Biology 14:44.

Svenson, G. J. and M. F. Whiting. 2009. Reconstructing the origins of praying mantises (Dictyoptera, Mantodea): the roles of Gondwanan vicariance and morphological convergence. Cladistics 25:468–514.
